# The age-related deceleration of clonal haematopoiesis in 420,000 healthy adults

**DOI:** 10.1101/2023.12.21.572706

**Authors:** Hamish A J MacGregor, Duc Tran, Kelly L Bolton, Douglas F Easton, Jamie R Blundell

**Affiliations:** Early Cancer Institute, University of Cambridge, United Kingdom; Centre for Cancer Genetic Epidemiology, Department of Public Health and Primary Care, University of Cambridge, United Kingdom; Division of Oncology, Department of Medicine, Washington University School of Medicine, St Louis, Missouri, United States

## Abstract

Somatic mutations acquired in haematopoietic stem cells can drive clonal expansions, a phenomenon known as clonal haematopoiesis (CH). CH has been associated with higher risk of haematological cancer, cardiovascular disease and lower life expectancy and is strongly age-dependent, becoming almost ubiquitous in older adults. However, the dynamics of CH are not yet fully understood, and an accurate quantitative model of clonal expansion in healthy people could inform strategies for risk stratification and early detection of haematological malignancy. Here, we analyse the age-dependence of the variant allele frequency (VAF) distributions of CH driver variants from blood-derived whole exomes in 420,000 cancer-free UK Biobank participants. Across a number of CH driver genes (including *DNMT3A, TP53* and *TET2*) we find evidence of a substantial deceleration of clonal expansion with age, while other drivers (including *SRSF2, SF3B1*, and *IDH2*) display more consistent growth throughout life. We find that this deceleration occurs consistently across the population and happens mostly before the age of 40. Using evolutionary models of clonal dynamics we assess the validity of alternative mechanisms of clonal deceleration. We find that variation in fitness among individuals (e.g. due to genetic susceptibility) is not sufficient to explain the deceleration. We also consider a clonal competition model in which clonal expansion later in life is inhibited by interference with a background of other expanding clones, and find this model inconsistent with the data given established estimates of the prevalence of hidden selection in blood. Our results imply that the intrinsic selection landscape in blood is substantially different in early life compared with middle age.

## Introduction

The acquisition and clonal expansion of somatic mutations is a ubiquitous feature of human ageing, observed across a range of tissues ^1–6^. In blood, this phenomenon, termed clonal haematopoiesis (CH), causes dramatic changes to the clonal diversity of haematopoietic stem cells (HSCs) in later life and is associated with higher risk of haematological cancers ^4,5^, cardiovascular disease ^7,8^, and lower life expectancy ^4^. CH is rare in young adults but the prevalence increases with age and it is observed in almost all adults by age 65^6^. A rapid collapse in clonal diversity is often observed around the age of 70^9^. However, we do not yet have a full quantitative understanding of the factors that shape this age dependence.

A simple model of CH in which driver mutations occur at a constant average rate and expand exponentially has been shown to capture key features of the variant allele frequency (VAF) distributions of clones, enabling one to estimate the fitness effects conferred by key CH-associated genes and mutation ‘hotspots’ ^10,11^. However, longitudinal studies of CH have yielded conflicting estimates of fitness, suggesting that clonal fitness may vary over a lifetime. The growth of driver variants in some genes (e.g. *DNMT3A, TP53*) has been suggested to decelerate with age, while other drivers (e.g. *SRSF2, SF3B1*) appear to expand more quickly in later life ^12^. Since most existing estimates of clonal expansion rates are derived from elderly people, little is known about the timing of age-related changes. The extent to which agerelated changes in fitness could be linked to variation between individuals, competition between clones in the HSC pool or age-related changes in HSC biology remains unknown.

UK Biobank (UKB) is a cohort of more than 420,000 individuals aged 37-73 at recruitment. Whole exome sequencing (WES) of blood samples from the cohort provides an opportunity to characterise CH at genomic hotspots across a large, healthy population spanning more than three decades. Here we use UKB to characterise VAF distributions of common CH driver mutations in people of different ages and investigate quantitatively how the dynamics of CH change with age. We observe that for many CH drivers there is a fundamental inconsistency between the growth rates required to explain the clones size distribution observed in the youngest age-group in UKB and the growth rates required to explain how the clone size distribution evolves between the youngest and oldest age-groups. By exploring mathematical models of clonal dynamics we find evidence for population-wide agerelated deceleration of CH that suggests that the selection landscape in blood in the first four decades of life is substantially different than in middle age.

## Results

### Widespread age-related deceleration of CH

Somatic clone sizes are determined by when variants arise and how quickly clones subsequently grow. Existing work has shown that somatic mutations in many tissues, including blood, are acquired at a constant rate throughout life, ^13^, and that the variation in clone sizes in CH is consistent with the principle that CH driver variants confer a consistent fitness advantage to cells per year ^10,14^, without the need to explicitly account for the spatial structure of the HSC niche or external selective pressures. However, emerging evidence suggests that the selection landscape in CH may change over individuals’ lifetimes. Clonal growth rates from longitudinal samples are frequently observed to be lower on average than those determined cross-sectionally, and some clones have growth rates too slow for them to have achieved their observed size within the lifespan of the individual ^9,12^, suggesting that deceleration of clonal expansion may be widespread in later life.

To try and reconcile these observations, we examined the VAF distributions across 424,089 individuals aged 37-73 from the UK Biobank exome sequencing dataset. We focussed our initial analysis on 18 common ‘hotspot’ variants for which sufficient data existed to analyse VAF spectra across a range of ages, excluding individuals with a history of cancer or those with multiple CH drivers (Methods). We applied an evolutionary framework we had previously developed ^10^ to estimate the fitness effect (*s*) associated with each variant by fitting the model to the VAF distribution observed across all ages (Fig. 1a), under the assumption of a constant expansion rate. The VAF distribution predicted by the model is in close agreement with the data, and the fitness effects and mutation rates inferred are broadly consistent with previous estimates (Fig. 1b, Supplementary Fig. S14, S15).

**Fig. 1.**
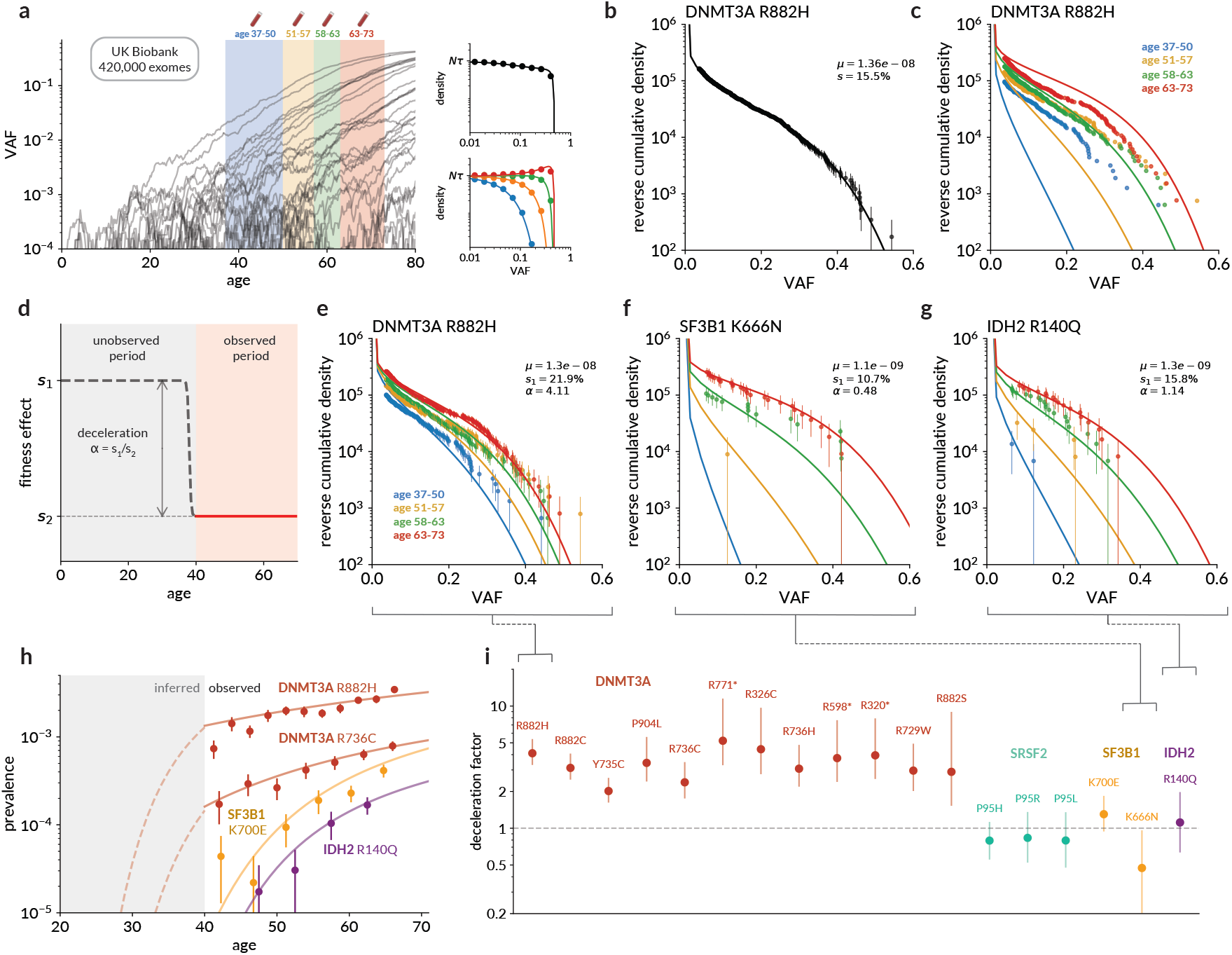
Diverse age-related dynamics across 18 CH hotspots in the UK Biobank. **(a)** Clones containing CH variants expand over individuals’ lifetimes (left, grey lines) and are sampled at single timepoints across a range of ages (simulated data). An evolutionary model predicts the VAF distributions of clones across a large cohort both overall (top right) and segregated by age (bottom right). **(b)** Cumulative density of *DNMT3A* R882H variants in UK Biobank (points) compared to expected density based on a constant growth rate model (line), with mutation rate *µ* (per cell per year) and fitness effect *s* (% per year) estimated using least-squares minimisation. Error bars were derived assuming counts were drawn from a Poisson distribution (*±* 1 s.e.). **(c)** Cumulative densities from (b) stratified into quartiles by age, each compared to its respective density predicted by the constant growth model. Two-phase model of growth, with deceleration factor *α* = *s*_1_*/s*_2_. **(e-g)** Cumulative VAF distributions for three hotspot variants: *DNMT3A* R882H (e); *SF3B1* K666N (f); *IDH2* R140Q (g), compared with expected density based on the two-phase model with parameters inferred using a maximum-likelihood approach. **(h)** Total prevalence of selected CH hotspot variants with age (points) compared with predictions from two-phase model (lines). Solid lines represent consistent exponential expansion at the late-life rate *s*_2_, dashed lines represent inferred early-life prevalence at the early-life rate *s*_1_ before age 40. **(i)** Estimated deceleration factors (*α*) for 18 common CH hotspot variants, with 95% confidence intervals derived from profile likelihoods.

However, stratifying the data by age uncovers a diversity of age-dependent growth rates. For several variants (e.g. *DNMT3A* R882H, Fig. 1c), the youngest UKB participants carry a substantially larger number of CH clones than would be predicted if clonal growth was consistent throughout life. These clones are also at much higher VAF than would be expected: some of the youngest individuals carry variants that have nearly fully swept the blood, which would require growth rates in excess of existing estimates. Conversely, the evolution of the VAF distributions between ages 40 and 70 years appears much slower than expected. This strongly suggests that the rate of clonal expansion driven by these variants varies over a lifetime.

### Quantifying age-dependent clonal dynamics

To quantify this age-dependent effect across multiple hotspots, we adopted a two-phase model of clonal dynamics in which clones grow at a constant rate *s*_1_ until age 40 years, and at a different rate *s*_2_ thereafter, with these rates consistent across the population (Fig. 1d). Very few UK Biobank participants were younger than 40 years at recruitment, so it is not possible to model the dynamics at younger ages: *s*_1_ simply represents an average growth rate in early life determined by the prevalence of variants at age 40. While the age-dependence of growth rates is likely to be more complex than this, this heuristic model can nonetheless be informative – for example, *s*_1_ establishes a lower bound on the early-life clonal growth rate, since if clones decelerate sooner than age 40 they must have expanded faster up to that point.

To fit the model, we developed a new maximum-likelihood approach which explicitly accounts for both the age and clone size for each individual (Supplementary Note 2D). Our method adjusts for the uncertainty in VAF introduced by sequencing at finite depth, an important source of additional dispersion in relatively low-depth sequencing data such as the UKB exomes (Supplementary Fig. S4).

We applied the two-phase model to each of the 18 CH hotspots, independently inferring the mutation rates (*µ*) and ‘deceleration factor’ (fold-change in fitness) *α* = *s*_1_*/s*_2_ for each variant. The model was able to capture the observed VAF distributions accurately (Fig. 1e-g, Supplementary Fig. S16). The age-specific prevalence was generally consistent with exponential growth over the 40-70 years age range, implying strongly that any deceleration occurs mostly before 40 years (Fig. 1h, Supplementary Fig. S16). In particular, for all 12 hotspots in *DNMT3A*, the two-stage model was a significant improvement (LR test: *p*< 5 × 10^−4^ for all DNMT3A variants), with each hotspot showing a substantial deceleration. Inferred values of the deceleration factor *α* were consistent across *DNMT3A*, between 2- and 5-fold (early life fitness effects 18-22%, later life 4-8%).

Expansion rates in the other genes however, the spliceosome genes *SRSF2* and *SF3B1* and the hotspot *IDH2* R140Q, appeared not to decelerate. The data for three common *SRSF2* P95 variants were consistent with the same selection parameters *s*, with the differences in prevalence caused by different mutation rates to each variant. The VAF distributions for these three variants, along with *SF3B1* K700E and *IDH2* R140Q, appeared consistent with a constant lifetime growth rate. *SF3B1* K666N was only seen in one individual under 58, and showed evidence of driving faster growth in later life (*p* = 0.037), albeit with less confidence than for the decelerating DNMT3A variants.

### Exome-wide CH dynamics

We next sought to characterise lifelong clonal dynamics across a wider range of CH driver genes. We called CH across the whole exome in the same UK Biobank participants using a set of bespoke filters (Methods and Supplementary Note 1B) and identified fourteen genes where at least 50 cancer-free individuals carried putative somatic clones, CH prevalence was associated with age at *p*< 0.05 and sequencing coverage was sufficient (Methods). (Fig. 2a). We aggregated data for all variants in each gene as a single variant ‘burden’, fitted the two-stage evolutionary model described above and estimated the total mutation rate to CH drivers and the early- and late-life fitness parameters for each gene. For genes where one or two hotspots (n > 50) dominated the CH spectrum (*MYD88, SRSF2*), only the hotspots were included and each was analysed separately. For *SRSF2*, these hotspots had already been included in the above analysis so they were not analysed again.

**Fig. 2.**
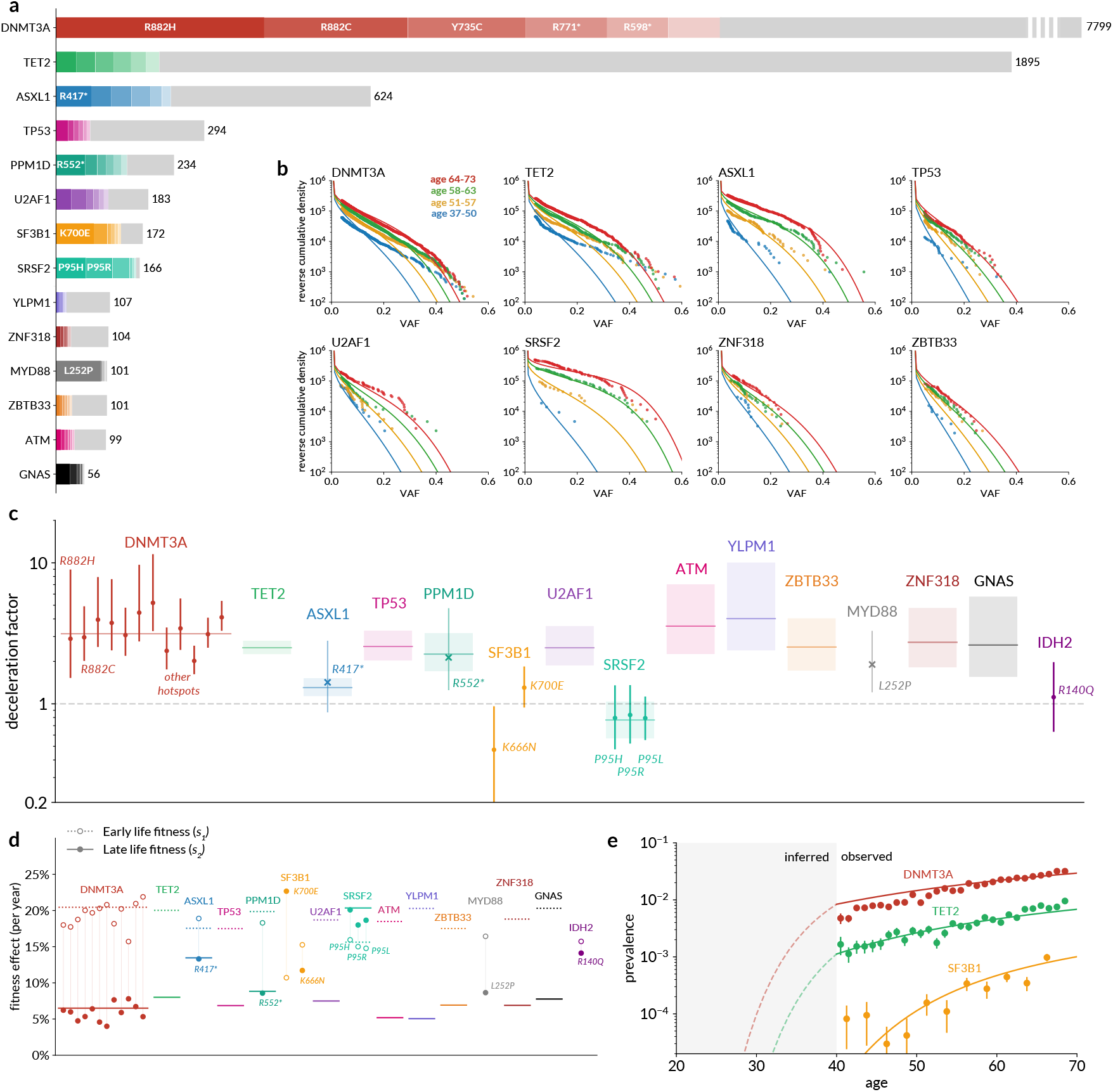
Clonal deceleration across the exome. **(a)** Number of putative CH variants called in each gene in the UK Biobank in healthy individuals. Ordered by most common variant in each gene, with the six most prevalent variants coloured. Genes are only shown if there are least 50 putative CH carriers in UKB and CH prevalence associated with age at *p*< 0.05. **(b)** Cumulative density of CH variants in selected genes in the UK Biobank (points) stratified by age, compared with expected density based on a two-stage evolutionary model. **(c)** Deceleration factors for all CH genes analysed (including the *SRSF2* and *IDH2* hotspots from the previous analysis). Horizontal lines represent estimated deceleration factor of variants aggregated by gene, showing 95% CI (shaded). Points represent individual hotspots, either from the original hotspot analysis (circles) or additional hotspots (n > 50) identified from the exome-wide calls (crosses). **(d)** Fitness effects (% per year) estimated for CH driver genes and hotspots using the two-stage model, both before the age of 40 (dotted lines, hollow circles) and afterwards (solid lines, shaded circles). Gene-wide estimates are shown as lines, hotspot-based estimates shown as circles. **(e)** Overall prevalence of CH variants in *DNMT3A, TET2* and *SF3B1* in the UK Biobank (points) compared with predicted prevalence from a two-stage evolutionary model.

Of the fifteen genes included in total, twelve displayed significant evidence of deceleration (*p*< 2.1 × 10^−4^), with only *SRSF2, SF3B1* (K700E and K666N) and *IDH2* (R140Q) consistent with consistent growth over a lifetime (*p>* 0.05). VAF distributions were broadly consistent with predictions from the two-stage model (Fig. 2b, Supplementary Figs. S16, S17). The magnitude of the deceleration in most genes was similar to *DNMT3A*, with deceleration factors *–* between 2 and 4 (Fig. 2c). An exception to this was *ASXL1*, where the deceleration was much less pronounced (*α* = 1.3, 95% CI 1.1-1.5) and the late-life growth rate remained relatively high (13% per year). For the genes exhibiting *DNMT3A*-like deceleration, expansion rates after age 40 were estimated to be 5-10% per year, with inferred early-life growth rates of approximately 20% per year (Fig. 2d). The deceleration of *DNMT3A* on aggregate was consistent with the behaviour of the twelve previously analysed *DNMT3A* hotspots, although the fit to the gene-wide VAF distribution was visibly poorer, likely as a result of aggregating a large number of drivers with varying fitness effects. A similar effect may be visible in TET2.

*TET2* has been observed to ‘overtake’ *DNMT3A* as the most prevalent CH driver gene in the over-75s ^12^. In UK Biobank, we found that *DNMT3A* was the most prevalent variant over the whole age range (Fig. 2e). While we observed a higher late-life growth rate for *TET2*-than *DNMT3A*-variant clones, *TET2*-variant clones presented clear evidence of early-life deceleration and a relatively slow expansion rate in later life, suggesting that its general behaviour is more similar to the majority of CH driver genes than to the late, fast-growing splicing genes.

### Fitness variation alone cannot explain age deceleration

To investigate how the underlying dynamics of the haematopoietic system might give rise to decelerating clones across the exome, we adapted our evolutionary model to explore two potential explanations: variation of fitness between individuals and clonal competition.

Longitudinal evidence suggests that clones with the same driver variant can expand at substantially different rates in different individuals ^12,15,16^. While the mechanisms underlying this diversity are not fully understood, genome-wide association studies (GWAS) have identified loci associated with prevalence of CH ^17–21^ suggesting fitness variation is partly driven by genetic susceptibility. To test whether variation in fitness among individuals could explain the apparent age-related deceleration of CH, we modified the existing model to draw fitness effects for each individual from a distribution of fitness effects (DFE). Fitness effects were drawn from a Gaussian distribution with mean 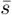 and standard deviation *σ* for each CH driver, which tended to a constant-fitness model as *σ* → 0 (Fig. 3a). We verified that introducing a DFE can cause changes in the VAF distribution that resemble deceleration, even if each individual’s fitness remains constant throughout life (Fig. 3b-c), Supplementary Note. 3B.1).

**Fig. 3.**
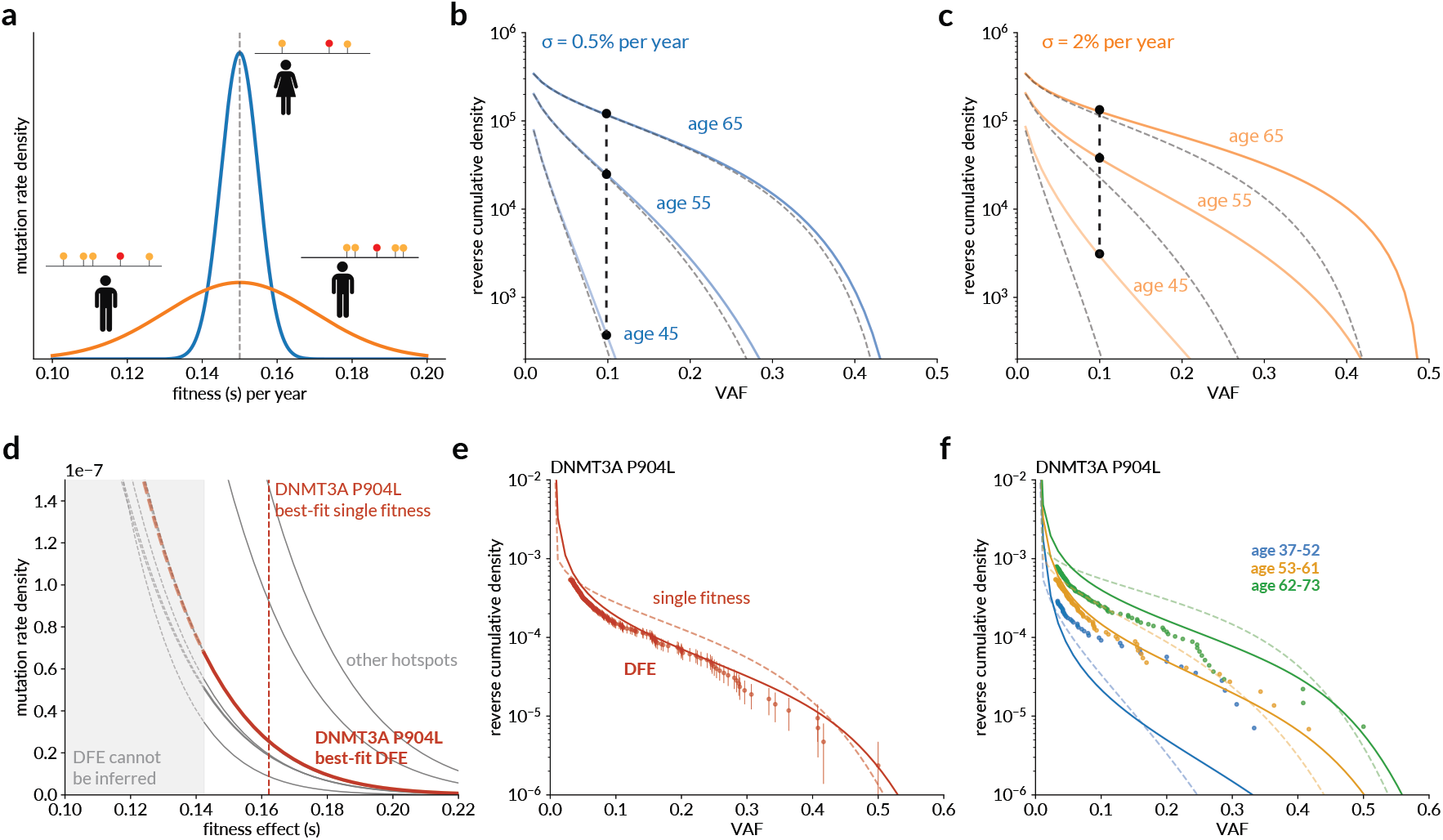
Variation in fitness among individuals with the same driver variant. **(a)** Schematic: individuals carrying the same CH driver variant, but with different germline susceptibility, experience different fitness effects drawn from a Gaussian distribution. Two examples shown: 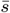, *σ* = 0.5%/yr (blue), 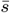, *σ* = 2%/yr (orange). **(b-c)** Reverse cumulative VAF distributions predicted by the narrower **(b)** and wider **(c)** DFEs, illustrating that a wider DFE produces an apparent deceleration in the VAF distributions (black points and line). Grey dashed lines show the VAF distribution for *σ* = 0 in each case. **(d)** Best-fit DFEs for the decelerating hotspots in UK Biobank (grey lines), with *DNMT3A* P905L highlighted as an example (maroon). Vertical dashed line shows the best-fit single fitness effect to *DNMT3A* P905L (*s* = 16.2%/yr). The shaded region and dashed lines approximately indicate the region where clones are insufficiently fast-growing to be observable in most UKB participants, and so the DFE in this region has no effect on predictions. **(e)** Reverse cumulative VAF distribution in UK Biobank for *DNMT3A* P905L (points) compared with prediction from best-fit DFE model (solid line) and the best-fit single fitness (*σ* = 0) (dashed line). **(f)** As in (e), stratified into three equal age cohorts. To allow comparison between different models, densities in (e) and (f) are not divided by mutation rate.

We applied this model to the decelerating CH drivers in UK Biobank to assess whether the deceleration observed could be accounted for by a distribution of fitness effects. We limited our analysis to individual hotspots since we would expect *a priori* that fitness would be more consistent at a single position across individuals. We estimated *µ*, 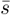 and *σ* for each of the twelve decelerating hotspots (all in *DNMT3A*) using a maximum-likelihood approach (Fig. 3d, Supplementary Fig. S18). Allowing a distribution of fitness effects improved the fit to the VAF distribution for each variant (*p*< 1.6 × 10^−4^) (Fig. 3e). However, despite the best-fit DFEs showing considerable variation in fitness (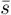 close to zero, *σ* ∼ 5% per year) the predicted deceleration remained notably weaker than observed in UKB. This suggests that fitness variation alone is not sufficient to explain the observed deceleration (Fig. 3f). The best fits to the data were achieved with monotonically decreasing DFEs, contrary to what might be expected if genetic susceptibility was the source of variation. However, our power to estimate the DFE at lower fitnesses is limited by the lower VAF threshold in the UKB exome data (3%), as variants with low fitness are very unlikely reach detectable levels in UK Biobank (Supplementary Note 3B.2).

### Competition-mediated deceleration requires very high levels of hidden selection

The genetic makeup of the blood changes with age, from a ‘polyclonal’ landscape comprising a large number of small, unique clones in early life to an ‘oligoclonal’ landscape dominated by a small number of highly expanded clones in later life ^9^. Competition between clones is a natural consequence of such a transition, and has been observed directly in individuals in the years prior to a diagnosis of acute myeloid leukaemia ^22^. If multiple clones of varying fitness reach a high VAF in the same individual, this could could slow, or even reverse, the clonal expansions of less fit clones later in life. A number of studies have shown that 60-80% of clonal expansions in blood do not involve alterations in currently known cancer driver genes ^9,14,23^, suggesting that deceleration could be due to competition from fitter clones with unobserved ‘hidden’ drivers.

We derived a model in which clones driven by such ‘hidden’ drivers can arise and expand alongside clones containing known CH drivers (Fig. 4a, Supplementary Note 3C) and estimated the parameters for the hidden drivers required to produce the deceleration. Both the observed and background clones were assumed to arise at a constant mutation rate and have constant fitness effects, with any age-dependent behaviour emergent from the competition between the two. We analysed all genes that had displayed significant evidence of deceleration, with the DNMT3A hotspots treated separately.

**Fig. 4.**
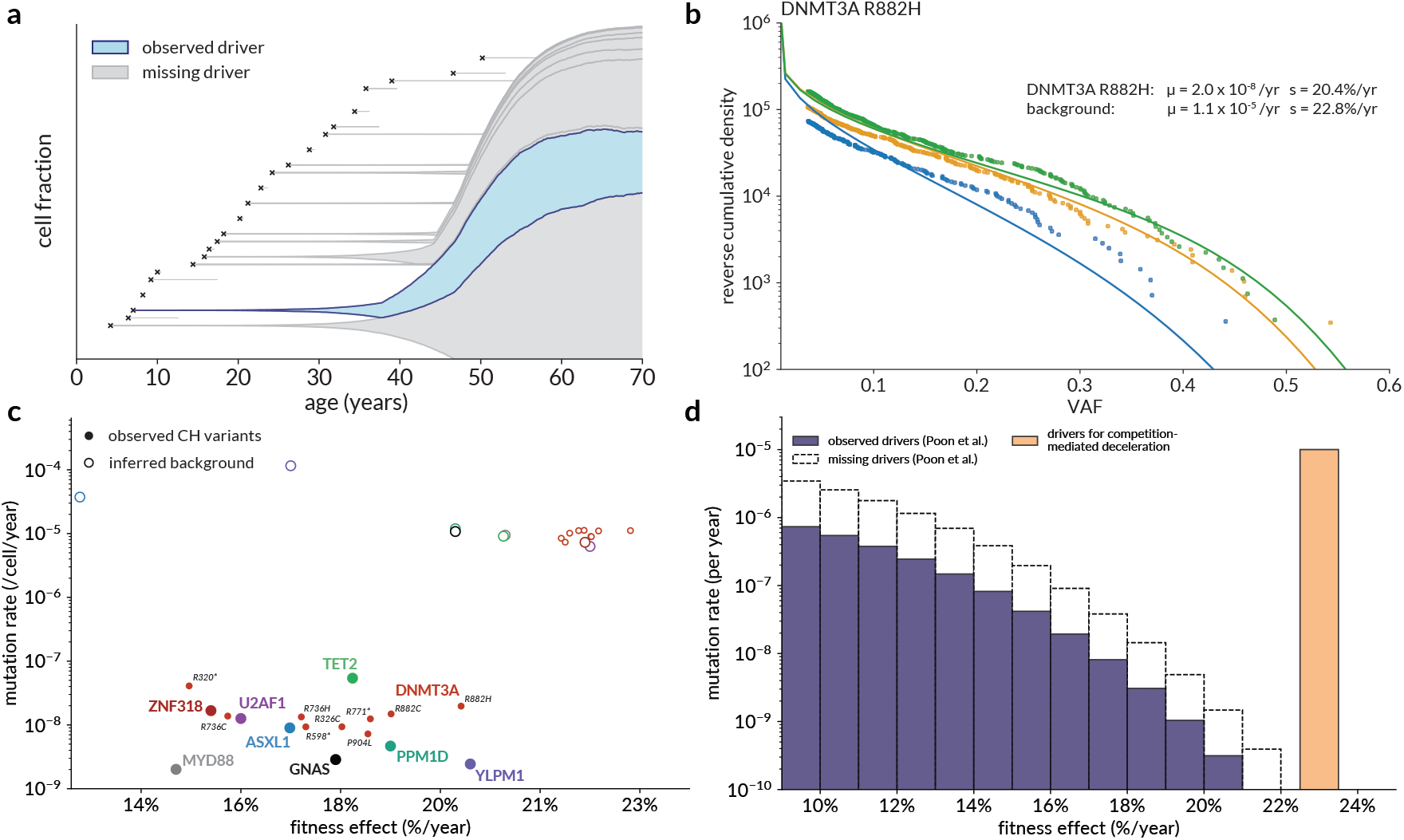
Competition-mediated clonal deceleration. **(a)** Simulated example of clonal expansion constrained by competition: a clone carrying an observed CH driver (*s* = 20% /year, blue) arises at a young age alongside unseen ‘background’ drivers of comparable fitness (*s* = 23%, grey) arising at a rate *µ* =1 × 10^−5^ per year. By middle age, the growth of the observed clone is constrained by the unseen expanding clones. **(b)** Cumulative density of UKB CH variants at the *DNMT3A* R882H hotspot (points) stratified by age, compared with expected density based on a clonal competition model with maximum-likelihood-fitted parameters. **(c)** Estimated mutation rates and fitness effects for decelerating variants under a clonal competition model. Solid circles represent *µ* and *s* for observed variants (aggregated by gene except for *DNMT3A*), and hollow circles represent estimated *µ* and *s* for hidden ‘background’ drivers required to produce deceleration. **(d)** Comparison of the levels of hidden selection required to produce competition-mediated deceleration in middle age (orange) and estimates by Poon et al. ^14^ of the distribution of fitness effects of observed (purple) and ‘missing’ (white) CH drivers, derived from the distribution of synonymous hitch-hikers.

For each of these genes and hotspots, we estimated the mutation rates and fitness effects for the CH hotspots and the unobserved background that would be required to obtain a good fit to the UKB data (Fig. 4b). The estimated fitness of the (unobserved) background clones was 20-23%, as expected slightly greater than the observed driver variants being analysed. The mutation rate (*µ*) of the background clones was ∼ 10^−5^ per cell per year, markedly higher than all the observed CH drivers (Fig. 4c).

It is also possible to estimate the characteristics of the background required for clonal competition in middle age from a theoretical perspective. To do this, we derived estimates for the the mutation rate and fitness of ‘hidden’ drivers required so that the average individual has at least one expanded clone by middle-age (detailed derivation in Supplementary Note 3C). We found that to reach the observed prevalence, the mutation rate to high-fitness drivers must be of the order 1*/N*_*τ*_ (*N* = HSC population size), that is ∼ 10^−5^ per cell per year. Moreover, only large-VAF clones will produce competition. For a typical clone to have reached a substantial VAF by middle age its fitness must be at least 20% per year.

A background mutation rate of approximately 10^−5^ per cell per year to highly fit drivers is significantly larger than existing estimates of the total mutation rate to CH drivers (including ‘missing’ drivers), for example by Poon et al. ^14^ (Fig. 4d). Exact comparisons are sensitive to assumptions about distribution of fitness, but the levels of missing selection required to produce extensive clonal competition in middle age are clearly inconsistent with the prevalence of hitch-hiking synonymous variants in blood (Supplementary Fig. S12). Such a large mutation rate to hidden drivers also implies that a significant fraction of HSCs in an average healthy person at age 40-50 are members of expanded clones (Supplementary Fig. S13). This is not observed in phylogenetic analysis of HSCs ^9^, where this ‘collapse’ in somatic diversity is seen only around the age of 70. We conclude, therefore, that clonal competition is unlikely to be the primary factor underlying the deceleration of clones observed in UK Biobank.

## Discussion

By analysing the cross-sectional VAF spectra of established CH driver variants across ∼ 420, 000 UK Biobank participants, we demonstrated that clonal expansion does not occur evenly across a person’s lifetime. The overabundance of variants seen in young people, relative to the expected frequency under a constant selective advantage, indicates that clonal expansion slows with age, and that much of this deceleration occurs before age 40 years. This deceleration was seen across the majority of, though not all, CH driver genes, suggesting that clonal deceleration is a more general phenomenon in CH.

Our findings are consistent in a number of important ways with existing analysis of CH derived from longitudinal samples in later life ^12^. Where previous cross-sectional analysis have estimated the average fitness of clones over a lifetime (e.g. *DNMT3A* R882H: *s* = 15%/yr ^10^), we were able to estimate the fitness in middle age directly, which we found to be in close agreement with longitudinal estimates (e.g. *DNMT3A* R882H: cross-sectional late-life *s* = 5.3%/yr, longitudinal *s* = 5.5%/yr ^12^). Similarly, longitudinal analyses have identified clones that expand too slowly in later life to explain the observed VAF under an assumption of constant growth, again implying early-life deceleration. Since the UK Biobank includes participants as young as 40 and growth rates appear consistent over the age range studied, we are able to conclude that any deceleration occurs primarily before the age of 40. An important implication is that clonal expansion in the young may be faster than previously thought (e.g. >20%/yr for *DNMT3A*). This prediction could be tested directly by studying children and young people, as discussed further below.

Another finding in agreement with existing work is that clones driven by *SRSF2* and *SF3B1* variants expand faster than other drivers in later life. We found that for these genes (and the *IDH2* R140Q hotspot), the data were consistent with a relatively low mutation rate but constant fitness throughout life. This indicates that their fitness advantage during later life might result from maintaining steady growth while most other clones slow down, rather than late-life acceleration as others have suggested ^24^. Clones carrying *ASXL1* variants were also found to retain a relative fitness advantage later in life. Conversely, clones carrying *U2AF1* variants showed modest late-life growth rates, although other studies have reported fast expansion in elderly individuals ^12^.

The model of deceleration presented is simplistic but illustrates that a period of fast growth followed by a period of slower (but consistent) growth captures the observed features of both the VAF distributions and prevalence curves in the UK Biobank. The nature of the early period is open to interpretation: for example, rapid clonal expansion in childhood followed by a decline during adolescence would produce the same VAF distributions as the model presented here, with suitably adjusted parameters (Supplementary note 3A). The findings raise the prospect of a yet-unknown biological change in HSC dynamics occurring in young people.

We sought to investigate how the fundamental dynamics of HSCs might give rise to clonal deceleration using mathematical models. Some of the key features of clonal dynamics, such as the collapse in clonal diversity around age 70, can be shown to be emergent properties of a constant-growth model ^9^. However, we found that competition between clones growing at a constant rate is unlikely to be sufficient to explain the deceleration. We estimated the genome-wide mutation rate and fitness effect of CH drivers that would be required to produce pervasive clonal competition in middle age, finding that both are implausibly large compared to existing estimates (for instance, derived from hitch-hiking synonymous variants ^14^). This is largely a consequence of the fact that the deceleration appears to occur relatively early in life: an alternative perspective is that pervasive clonal competition is a feature of the collapse in diversity, and before this collapse occurs clonal competition is unlikely to play a major role at the population level.

The variation in clonal expansion rates between healthy individuals carrying the same CH driver remains relatively poorly characterised. A weakness of our single-timepoint approach is that the the effects of deceleration and fitness variation are mutually confounding and therefore difficult to quantify independently. We found that incorporating variation in fitness effects improved predictions of VAF distribution, consistent with evidence from longitudinal studies ^6,12,15,25^. However, we found that even an exaggerated degree of variation was unable to reproduce the extent of clonal deceleration observed in UK Biobank, suggesting that while variation is probably present, it is not the primary mechanism responsible for the apparent slowing of clonal growth. Longitudinal studies are likely better suited to addressing this issue, but age again may be an important factor: in older people with a clonal spectrum characterised by a few highly-expanded clones, clonal competition and multiple mutation are likely to increase observed variation in fitness. Since most existing longitudinal studies are in older people, the degree of inter-individual diversity in the 40-70 year age group is even less clear. Higher-VAF clones are associated with increased cancer risk ^4^, so clonal expansion rate at the individual level could be an important risk modifier: this is a key area for future study.

The mechanism underlying the observed deceleration remains unknown. Changes in stem cell numbers over time could produce decelerating dynamics, and existing work has identified decreasing stem cell division rates with age in colon, oesophagus and other non-blood tissues in humans ^26^. The HSC population in blood, however, has been estimated to be quite stable at *N*_*τ*_ ≈ 100, 000 from early childhood onwards, albeit with a relatively large uncertainty ^9,27^. One feature of our evolutionary model is that HSC population size is better estimated using low-VAF clones, so the limited sequencing depth of the UKB exomes constrains our ability to assess this issue. Other mechanisms have been proposed that could give rise to apparent age-related deceleration: for instance, that for some driver genes, decaying telomere length in expanding clones impedes cell division later in life ^24,28–30^.

A more complete model of healthy CH dynamics would be greatly facilitated if CH could be evaluated directly in individuals from early childhood to middle age. A sufficiently large and deeply-sequenced study (ideally with data at multiple time points) would enable many of the poorly-understood aspects of healthy CH to be explored in detail: clonal fitness in early life and the mechanisms of the deceleration, the effects of germline genetics and extrinsic environmental factors on fitness and mutation rate, and the role of early development and HSC population size.

## Methods

### Identifying CH in the UK Biobank

Exome sequencing of 450,000 people was performed on DNA from blood samples from participants in the UK Biobank, ^31,32^, a prospective cohort study of 500,000 middle-aged people in the UK. All UKB participants signed a written informed consent form at enrolment. Ethical approval was given by the North West Multi-Centre Research Ethics Committee (REC 21/NW/0157)

Individuals with unknown age at blood draw were excluded. Since some types of systemic cancer treatment have been shown to influence clonal expansion in blood ^33^, individuals with a history of cancer were excluded. This included self-reported cancers, but excluded basal cell carcinoma, rodent ulcer and pre-cancer of the cervix. After these filters were applied, a total of 424,089 individuals remained in the cohort.

To isolate as far as possible the ‘intrinsic’ fitness conferred by a single driver variant, individuals with more than one detectable CH mutation were excluded. Individuals with mosaic chromosomal alterations (mCA, called by Loh. et al ^34^) overlapping the variant locus were also excluded on the same grounds ^11^.

The exact approach to calling CH differed depending on whether we were interrogating only CH hotspots or the whole exome, owing to the varying prior probability of finding CH at different genomic positions.

### Variant calling at CH hotspots

Variants at established CH hotspots are much more likely to be acquired somatically than in the germline. Therefore, we took a more permissive approach to calling CH at these positions to maximise our ability to resolve trends in age dependence.

CH hotspots were called from the UK Biobank exome CRAM files using the DNANexus Research Analysis Platform (application 28126) ^35^. *samtools mpileup* (docker: *quay*.*io/biocontainers/samtools:1*.*12–hd5e65b6_0)*) was used to identify variants at 26 genomic positions in 11 genes, corresponding to common single nucleotide variants identified as potential drivers of CH by Watson et al. ^10^ (Supplementary table S1). Variants supported by fewer than 3 variant reads were excluded. Above this threshold, all variants were assumed to be somatic. Variants carried by fewer than 40 individuals in UKB were excluded. A low-VAF filter was applied for each variant to minimise the impact of false negatives at low VAF (Supplementary Note 1A, Supplementary Fig. S3). At the positions of the common hotspots *JAK2* V617F and *GNB1* K57E, average sequencing depth was too low for reliable parameter estimation, so these variants were excluded from the analysis. Variants were annotated using Ensembl variant effect predictor v111.0^36^. 18 hotspot variants in 4 genes passed all filters. Across all these variants, the total number of CH carriers was 3,804.

### Exome-wide CH calling

Exome-wide CH calling was performed on the UK Biobank CRAM files using the somatic variant callers Mutect2 and VarDictJava, along with a more extensive set of filters aiming at excluding artefacts and germline variants. This approach had been applied previously to a subset of the UKB cohort, and full details of the calling procedure are available in that publication ^37^. A summary of the calling method and filters are found in Supplementary Note 1B. Genes with fewer than 50 carriers of putative drivers (after trimming for false negatives as above) were excluded. Some genes showed a weak association between CH prevalence and age, suggesting the presence of artefacts, germline mutations or developmental mutations despite the above filters (Supplementary Note 1B). We retained only genes with a positive association between CH and age at *p*< 0.05 (logistic regression with sex as a covariate, Supp. Fig. S2). Three genes where low sequencing depth affected the reliability of parameter estimation (*GNB1, JAK2, CBL*) were also excluded. Variants were annotated using *Ensembl variant effect predictor v111*.*0* ^36^. In total, 14 genes passed all filters. In these genes, a total of 12,179 putative CH carriers were identified from a cohort of 422,882 individuals. The slight difference in cohort size was caused by the different versions of the UK Biobank data available at the respective times of the analysis.

### Mathematical modelling

The details of the mathematical models, parameter estimation methods and simulations used in this analysis are provided in the supplementary material. All simulations were carried out using custom Python scripts.

## Acknowledgements

We thank Arnold Levine, Daniel Fisher and Dan Landau for helpful discussions. We thank all members of the Blundell lab for input. This research has been conducted using the UKB Resource under application number 28126. JRB is funded by a UKRI Future Leaders Fellowship and by the CRUK Cambridge Cancer Centre. DFE is supported by the NHS in the East of England through the Clinical Academic Reserve. HAJM is supported by the International Alliance for Cancer Early Detection, an alliance between Cancer Research UK (C14478/A29329), Canary Center at Stanford University, the University of Cambridge, OHSU Knight Cancer Institute, University College London and the University of Manchester.

## Author Contributions

Simulations and mathematical models were developed by JRB and HAJM with input from DFE. CH hotspot variant calling was performed by HAJM with input from JRB and DFE. Exome-wide CH calling was planned and performed by DT and KLB, with some additional filters applied by HAJM with input from DFE and JRB. Manuscript was written by HAJM and JRB and edited by all authors.

