## Supplementary materials for "The age-related deceleration of clonal haematopoiesis in 420,000 healthy adults"

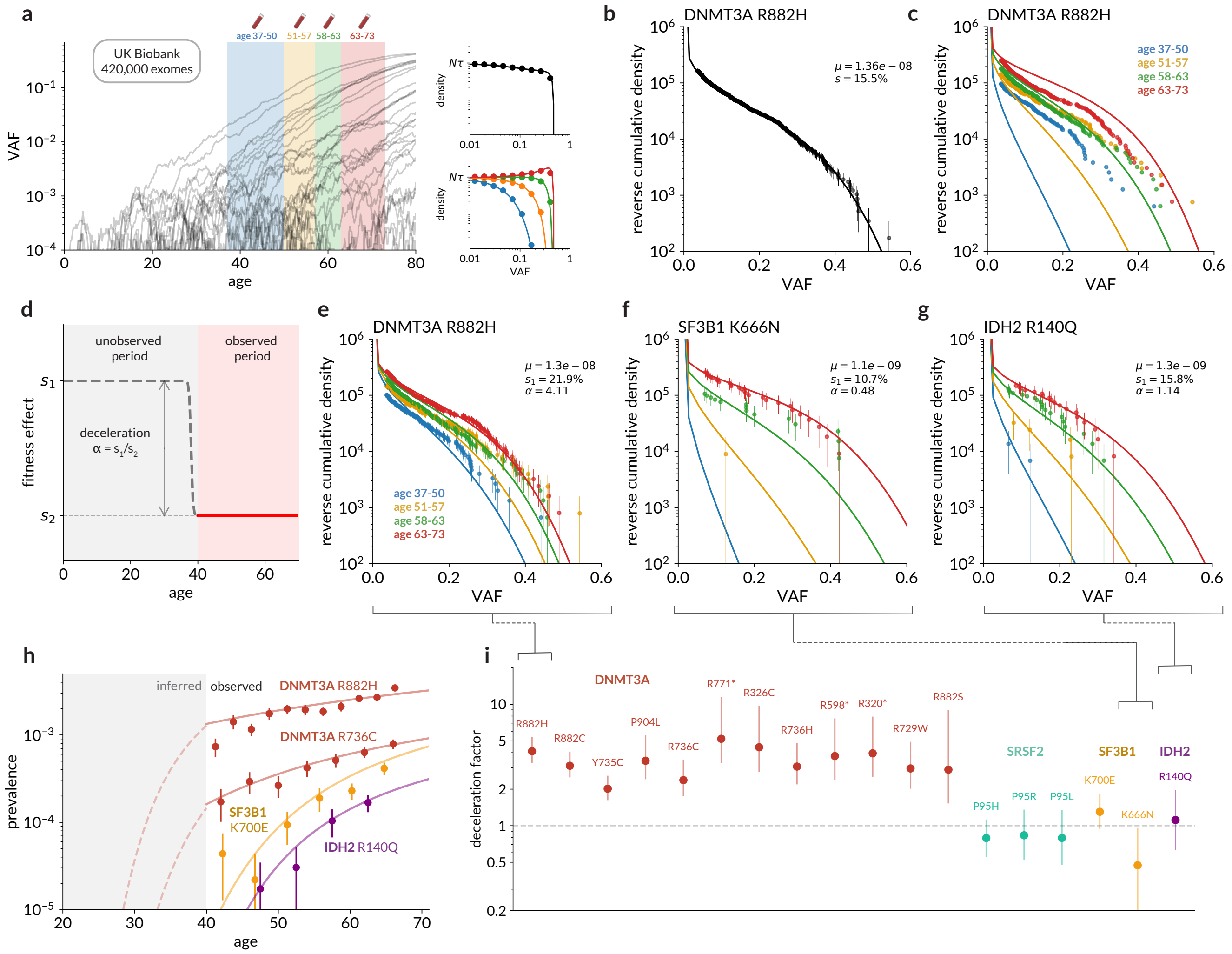

**Fig. 1. Diverse age-related dynamics across 18 CH hotspots in the UK Biobank**. **(a)** Clones containing CH variants expand over individuals' lifetimes (left, grey lines) and are sampled at single timepoints across a range of ages (simulated data). An evolutionary model predicts the VAF distributions of clones across a large cohort both overall (top right) and segregated by age (bottom right). **(b)** Cumulative density of *DNMT3A* R882H variants in UK Biobank (points) compared to expected density based on a constant growth rate model (line), with mutation rate $\mu$ (per cell per year) and fitness effect $s$ (% per year) estimated using least-squares minimisation. Error bars were derived assuming counts were drawn from a Poisson distribution ($\pm 1$ s.e.). **(c)** Cumulative densities from (b) stratified into quartiles by age, each compared to its respective density predicted by the constant growth model. **(d)** Two-phase model of growth, with deceleration factor $\alpha = s_1/s_2$. **(e-g)** Cumulative VAF distributions for three hotspot variants: *DNMT3A* R882H (e); *SF3B1* K666N (f); *IDH2* R140Q (g), compared with expected density based on the two-phase model with parameters inferred using a maximum-likelihood approach. **(h)** Total prevalence of selected CH hotspot variants with age (points) compared with predictions from two-phase model (lines). Solid lines represent consistent exponential expansion at the late-life rate $s_2$, dashed lines represent inferred early-life prevalence at the early-life rate $s_1$ before age 40. **(i)** Estimated deceleration factors ($\alpha$) for 18 common CH hotspot variants, with 95% confidence intervals derived from profile likelihoods.

Longitudinal evidence suggests that clones with the same driver variant can expand at substantially different rates in different individuals[12,15,16]. While the mechanisms under-

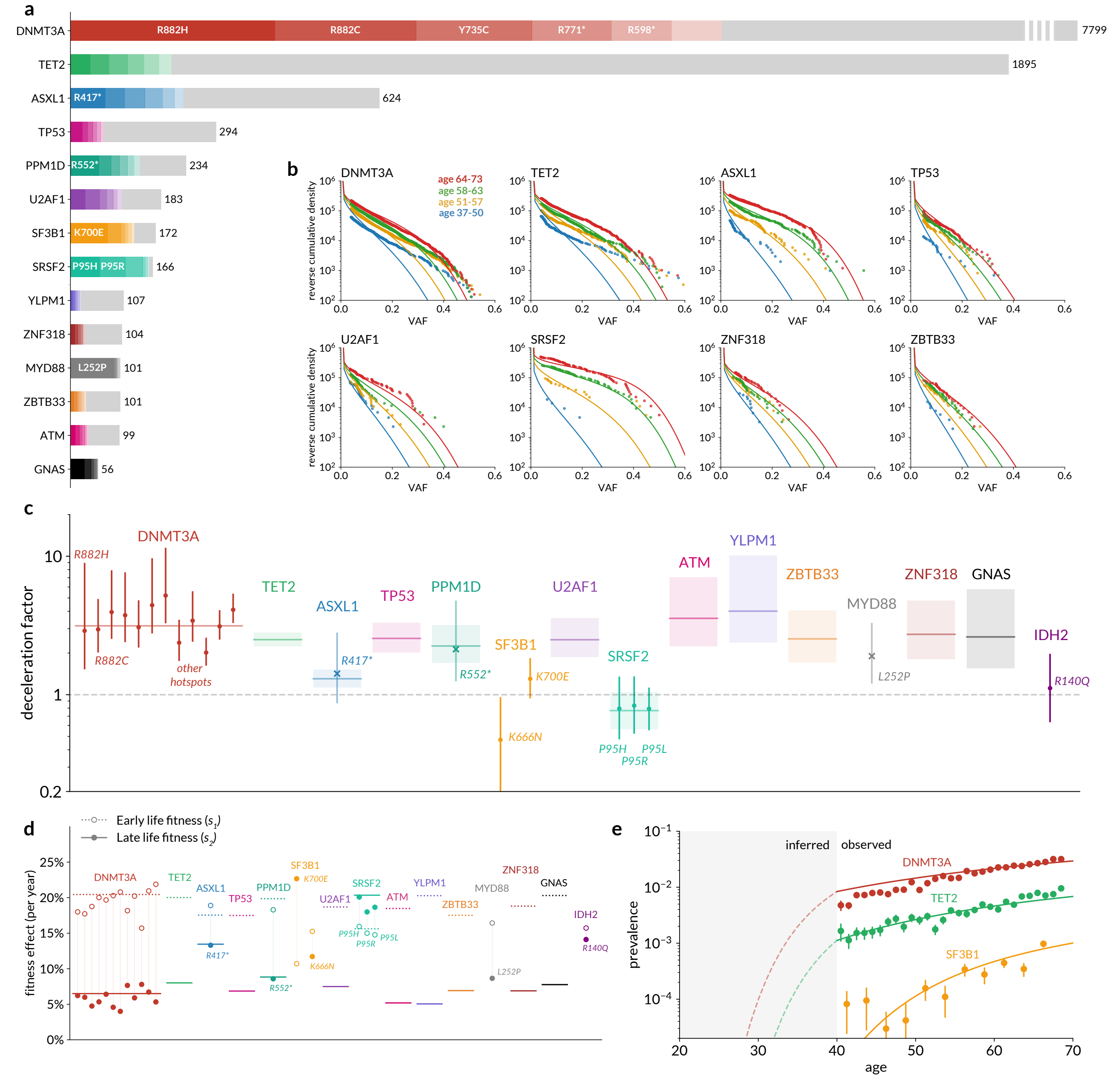


**Fig. 2. Clonal deceleration across the exome.** **(a)** Number of putative CH variants called in each gene in the UK Biobank in healthy individuals. Ordered by most common variant in each gene, with the six most prevalent variants coloured. Genes are only shown if there are least 50 putative CH carriers in UKB and CH prevalence associated with age at $p < 0.05$. **(b)** Cumulative density of CH variants in selected genes in the UK Biobank (points) stratified by age, compared with expected density based on a two-stage evolutionary model. **(c)** Deceleration factors for all CH genes analysed (including the *SRSF2* and *IDH2* hotspots from the previous analysis). Horizontal lines represent estimated deceleration factor of variants aggregated by gene, showing 95% CI (shaded). Points represent individual hotspots, either from the original hotspot analysis (circles) or additional hotspots (n > 50) identified from the exome-wide calls (crosses). **(d)** Fitness effects (% per year) estimated for CH driver genes and hotspots using the two-stage model, both before the age of 40 (dotted lines, hollow circles) and afterwards (solid lines, shaded circles). Gene-wide estimates are shown as lines, hotspot-based estimates shown as circles. **(e)** Overall prevalence of CH variants in *DNMT3A*, *TET2* and *SF3B1* in the UK Biobank (points) compared with predicted prevalence from a two-stage evolutionary model.

lying this diversity are not fully understood, genome-wide association studies (GWAS) have identified loci associated with prevalence of CH[17–21] suggesting fitness variation is partly driven by genetic susceptibility. To test whether variation in fitness among individuals could explain the apparent age-related deceleration of CH, we modified the existing model to draw fitness effects for each individual from a distribution of fitness effects (DFE). Fitness effects were drawn from a Gaussian distribution with mean $\bar{s}$ and standard deviation $\sigma$ for each CH driver, which tended to a constant-fitness model as $\sigma \to 0$ (Fig. 3a). We verified that introducing a DFE can cause changes in the VAF distribution that resemble deceleration, even if each individual's fitness remains constant throughout life (Fig. 3b-c), Supplementary Note. 3B.1).

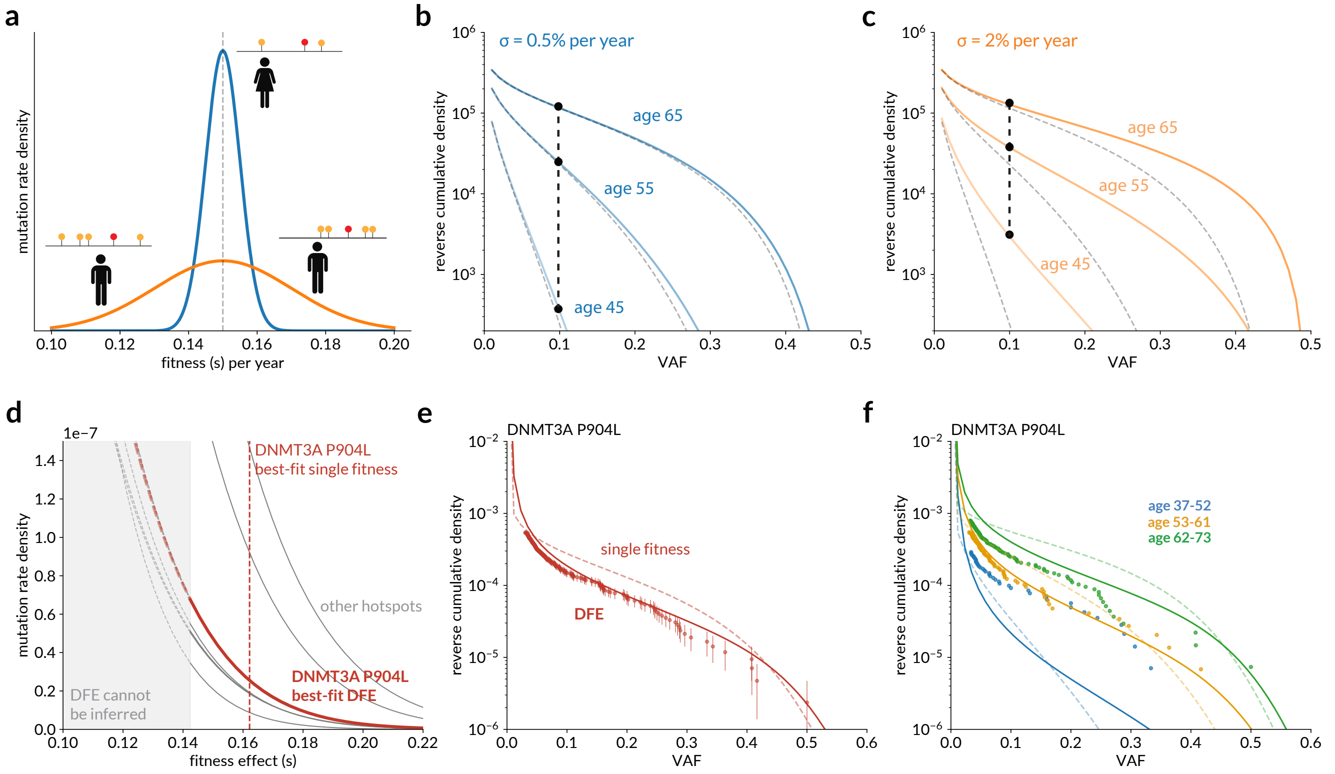


**Fig. 3. Variation in fitness among individuals with the same driver variant. (a)** Schematic: individuals carrying the same CH driver variant, but with different germline susceptibility, experience different fitness effects drawn from a Gaussian distribution. Two examples shown: $\bar{s} = 15\%$/year, $\sigma = 0.5\%$/yr (blue), $\bar{s} = 15\%$/yr, $\sigma = 2\%$/yr (orange). **(b-c)** Reverse cumulative VAF distributions predicted by the narrower **(b)** and wider **(c)** DFEs, illustrating that a wider DFE produces an apparent deceleration in the VAF distributions (black points and line). Grey dashed lines show the VAF distribution for $\sigma = 0$ in each case. **(d)** Best-fit DFEs for the decelerating hotspots in UK Biobank (grey lines), with *DNMT3A* P905L highlighted as an example (maroon). Vertical dashed line shows the best-fit single fitness effect to *DNMT3A* P905L ($s = 16.2\%$/yr). The shaded region and dashed lines approximately indicate the region where clones are insufficiently fast-growing to be observable in most UKB participants, and so the DFE in this region has no effect on predictions. **(e)** Reverse cumulative VAF distribution in UK Biobank for *DNMT3A* P905L (points) compared with prediction from best-fit DFE model (solid line) and the best-fit single fitness ($\sigma = 0$) (dashed line). **(f)** As in (e), stratified into three equal age cohorts. To allow comparison between different models, densities in (e) and (f) are not divided by mutation rate.

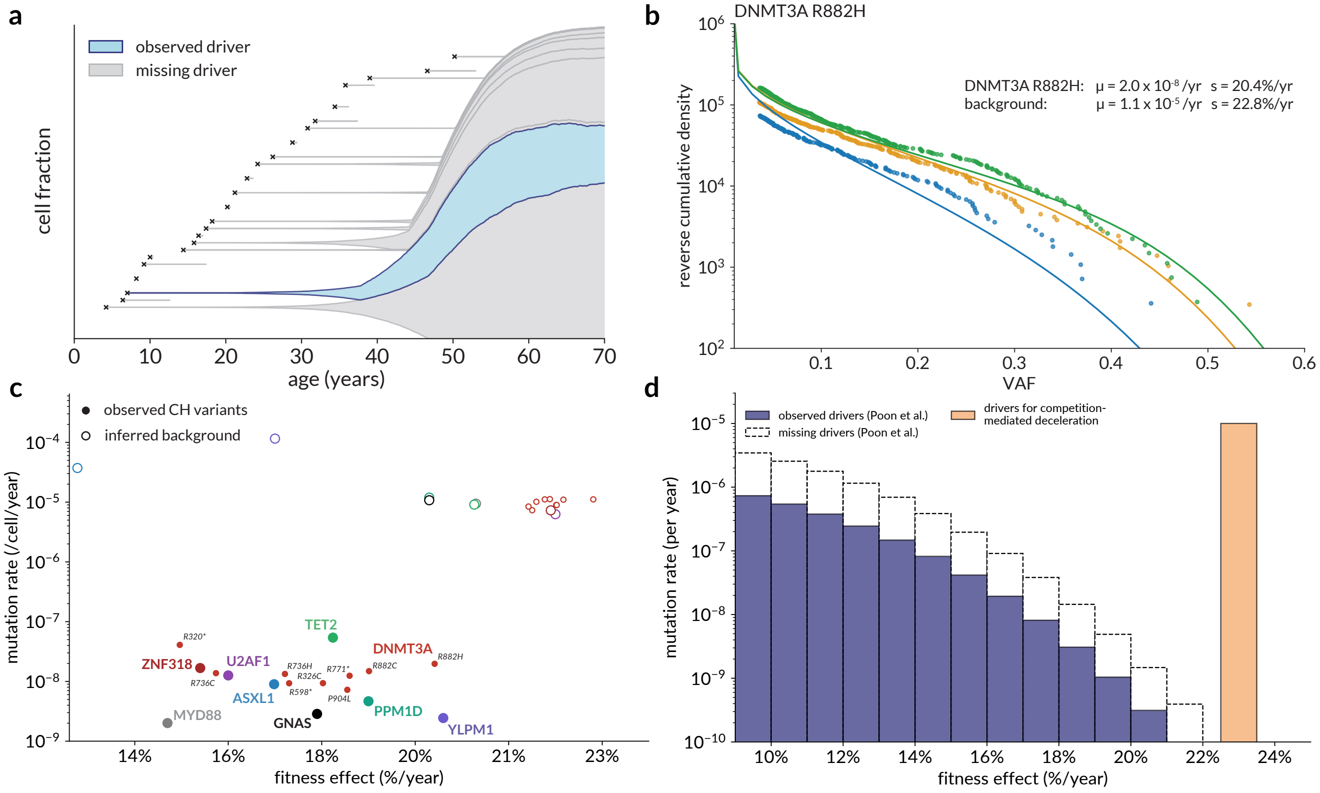


**Fig. 4. Competition-mediated clonal deceleration.** **(a)** Simulated example of clonal expansion constrained by competition: a clone carrying an observed CH driver ($s = 20\%$ /year, blue) arises at a young age alongside unseen 'background' drivers of comparable fitness ($s = 23\%$, grey) arising at a rate $\mu = 1 \times 10^{-5}$ per year. By middle age, the growth of the observed clone is constrained by the unseen expanding clones. **(b)** Cumulative density of UKB CH variants at the *DNMT3A* R882H hotspot (points) stratified by age, compared with expected density based on a clonal competition model with maximum-likelihood-fitted parameters. **(c)** Estimated mutation rates and fitness effects for decelerating variants under a clonal competition model. Solid circles represent $\mu$ and $s$ for observed variants (aggregated by gene except for *DNMT3A*), and hollow circles represent estimated $\mu$ and $s$ for hidden 'background' drivers required to produce deceleration. **(d)** Comparison of the levels of hidden selection required to produce competition-mediated deceleration in middle age (orange) and estimates by Poon et al.[14] of the distribution of fitness effects of observed (purple) and 'missing' (white) CH drivers, derived from the distribution of synonymous hitch-hikers.

### Supplementary Note 1: CH in the UK Biobank

**A. Hotspot variant calling and filtering.** Table S1 shows the 27 positions at which putative CH driver variants were called using *samtools mpileup* from the UK Biobank exome CRAM files. The variants were all identified as common CH drivers by Watson et al.[10]. Individuals with fewer than 3 variant reads or with more than one hotspot variant were excluded. Variants with overlapping mosaic chromosomal alterations (identified by Loh et al.[38] were also discarded. It was also necessary to impose an additional VAF threshold, since there is a region at low VAF where the variable sequencing depth (and random sampling error) means that a substantial fraction of variants are likely to be missed (false negatives). Following Watson et al., we imposed the VAF threshold for each variant at a point where the density began to decline (Fig. S3, Table S2). After all filters were applied, at least 40 carriers were required for the variant to be included in the final analysis. Two additional variants were excluded from the main analysis on the grounds that their mean sequencing depth was too low to generate reliable estimates of fitness and mutation rate using the maximum-likelihood methods described below: *JAK2* V617F (mean depth 24 reads) and *GNB1* K57E (mean depth 22 reads).

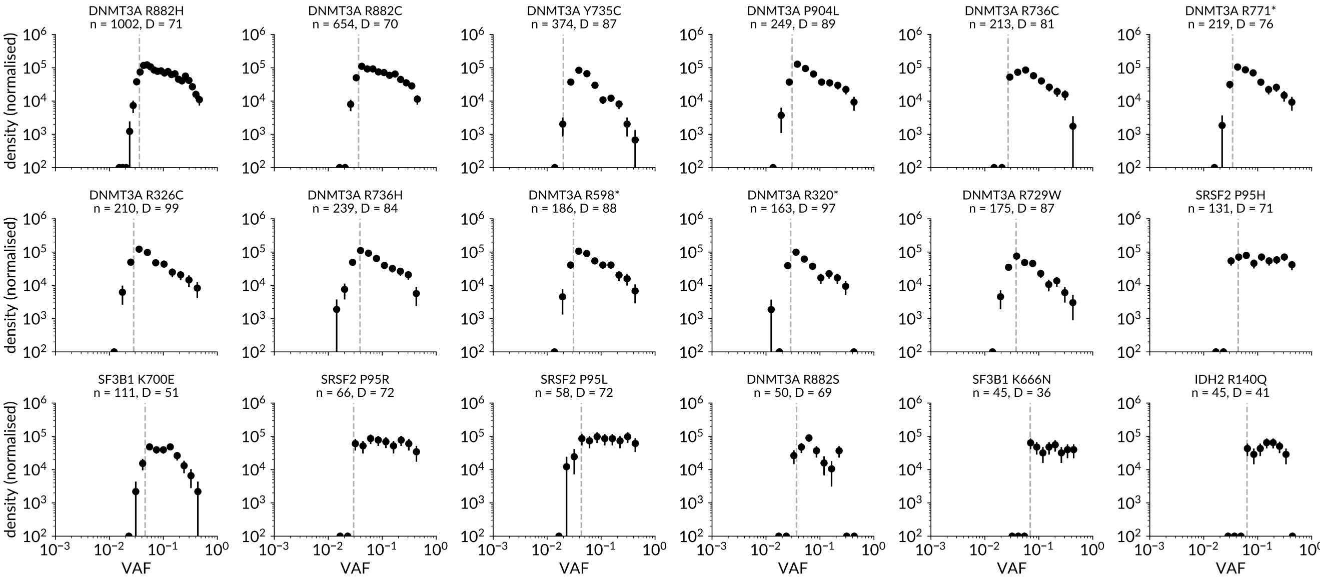


**Fig. S1. VAF distributions at CH hotspots:** Overall VAF distributions of CH variants at 18 hotspots, showing number of variant carriers $n$ and mean sequencing depth $D$. Vertical dashed line shows the lower VAF threshold imposed to exclude regions dominated by false negatives. Values of density are normalised vertically by inferred mutation rate from the two-stage model.

#### B. Exome-wide CH calling.

***B.1. Exome-wide calling from UKB CRAM files.*** Exome-wide calling was done using the procedure described in a recent publication by Tran et al. (2024)[37]. Full details are available in Supplementary Note 1 of that publication. In summary, variants were called from the UKB exome CRAM files using Mutect2[39] and VarDictJava[40]. Variants were retained if they had at least 2% VAF and were supported by at least one read on each of the forward and reverse strands. A panel of normals derived from UKB individuals was used to filter for sequencing error. Filters were applied to remove potential germline polymorphisms, including: comparison with the gnomAD database[41]; removal of variants with high VAF unless at somatic hotspots or previously reported as CH. For a variant to qualify as a putative CH driver, it also needed to satisfy at least one of a number of further conditions based on predicted functional effect or previous identification as CH (details in Tran et al.[37]).

***B.2. Association of CH with age.*** After the exome-wide calling pipeline had been applied, variants in a number of genes identified as putative CH drivers exhibited very weak or non-existent association between CH prevalence and age. Tellingly, CH with negligible age-association was most prevalent in genes with very long coding lengths such as *KMT2C*, *KMT2D* and *NF1*. This suggested that these variants were either acquired during early embryonic development and ‘hitch-hiked’ to high VAF (but conferred no significant clonal fitness in their own right) or were simply artefacts that had escaped the filters. We compared the VAF spectra with the distributions expected for developmental mutations (following Poon et al.[14], more detail in supplementary note 2F). The prevalence of CH observed in these genes was too large to be explained by neutral developmental hitch-hiking given current best estimates for mutation rates in development. CH in these genes is therefore likely to be dominated by sequencing artefacts. We performed a logistic regression using Python *statsmodels 0.14* (variant present/absent $\sim$ age + sex M/F), and excluded any gene that did not show positive age-association at a significance of at least $p < 0.05$ (Fig. S2).

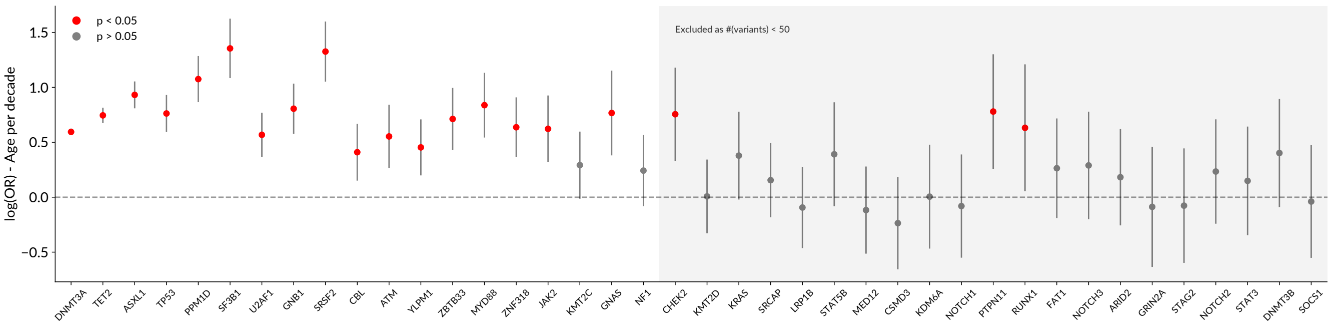


**Fig. S2. Association of CH with age:** log-odds ratios for the association between presence/absence of CH in each gene with age (by decade), with sex as a covariate. Error bars represent 95% confidence intervals. The grey shaded area represents genes which were excluded on the grounds that there were fewer than 50 CH carriers in the UK Biobank.

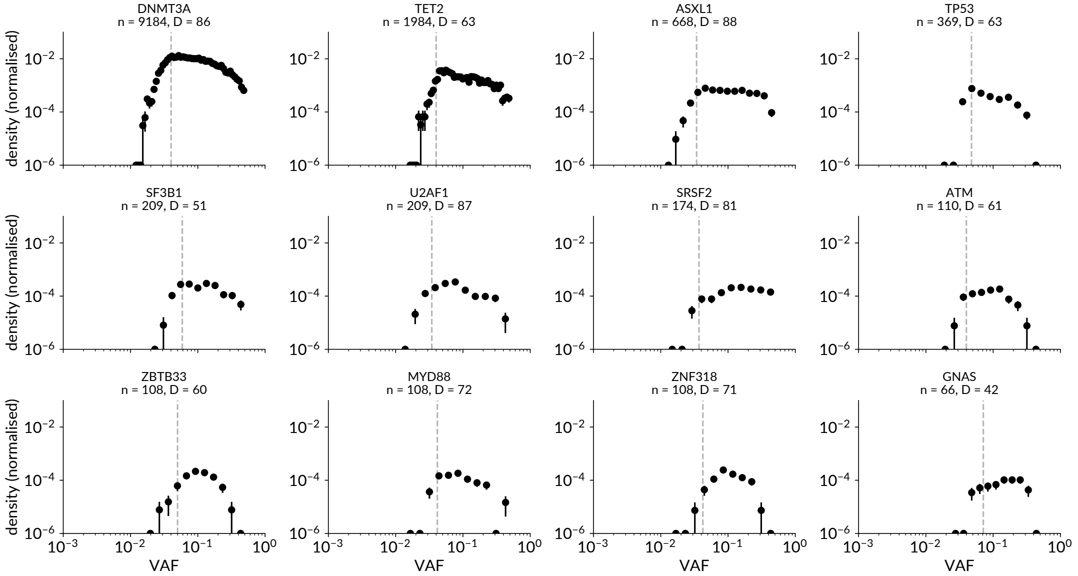


**Fig. S3. VAF distributions aggregated across CH genes:** Overall VAF distributions of CH variants in 14 driver genes, showing number of variant carriers $n$ and mean sequencing depth $D$. Vertical dashed line shows the lower VAF threshold imposed to exclude regions dominated by false negatives. Values of density are normalised vertically by inferred mutation rate from the two-stage model.

### Supplementary Note 2: Quantitative modelling of clonal haematopoiesis

**A. Evolutionary model of CH with constant growth rate.** The work presented here builds on the constant-growth model of CH developed by Watson et al. (2020)[10], and full details of the model and its derivation are found in that publication. In summary, a population of HSCs of size $N$ divides self-symmetrically with average division time $\tau$. Mutations enter this population stochastically at a constant rate $\mu$ per cell per year. A cell with a mutation experiences a constant (variant-specific) fitness effect $s$ which biases it towards self-renewal, such that an established clone of mutant cells grows exponentially at rate $s\%$ per year. From this framework it is possible to derive the expected distribution of clone sizes $n$ across a population of individuals at age $t$ years[10,42]:

$$\rho(n,t,\{a_i\})dn = n^{\theta-1}\frac{e^{-n/\tilde{n}(t)}}{\Gamma(\theta)\tilde{n}(t)^{\theta}}dn \tag{1}$$

$$\text{where} \quad \theta = N\tau\mu \quad \text{and} \quad \tilde{n}(t) = \frac{\exp(st)-1}{s\tau}$$

$\tilde{n}(t)$ can be interpreted as the characteristic scale of the largest clones in a person of a given age $t$, in units of number of cells. For the constant-growth model the parameters $\{a_i\} = \{\mu, s\}$; in the alternative growth models these are replaced by new parameters. A change of variables ($2f = \frac{n}{N+n}$) gives the expected distribution of VAF $f$ :

$$\rho(f,t,\{a_i\})df = \frac{(2N)^{\theta}}{f^{1-\theta}(1-2f)^{1+\theta}}\exp\left(\frac{-2Nf}{\tilde{n}(t)(1-2f)}\right)\frac{1}{\Gamma(\theta)\tilde{n}(t)^{\theta}}df \tag{2}$$

This density integrates to 1 over the interval $0 < f < 0.5$ (the maximum attainable VAF for a heterozygous mutant clone assuming that an identical second mutation does not occur on the other allele), meaning that it can be treated as a probability density function for finding a variant at VAF $f \to f + df$ in an individual of age $t$.

The ‘prevalence’ of a CH variant was defined as the total fraction of individuals in whom a given CH variant was observed, regardless of VAF. Predictions of prevalence were obtained by integrating the VAF distribution from the lower detection limit in UKB (imposed by trimming) to 0.5:

$$\text{Prev}(t, \{a_i\}) = \int_{f_0}^{0.5} \rho(f,t,\{a_i\}) df$$

Predictions for variant density and prevalence for a population with a range of ages were obtained using a weighted sum of single-age predictions, weighted by the distribution of sampling ages.

**B. HSC population size.** Our inferences rely on the value of the parameter $N\tau$, the product of number of self-renewing HSCs ($N$) and the self-symmetric division time ($\tau$). While these parameters are difficult to estimate independently, the product has been estimated independently by Lee-Six et al.[27] and Mitchell et al.[9] from single-cell phylogenetics and by Watson et al.[10] from single-timepoint VAF distributions. Estimates from these studies consistently suggest that $N\tau \approx 100{,}000$ cell-years. Mitchell et al. also show that $N\tau$ stabilises early in childhood and remains approximately constant into adulthood.

It is possible to estimate $N\tau$ from single-timepoint VAF spectra such as UK Biobank, as done in Watson et al.[10]. However, for two reasons we have chosen not to estimate $N\tau$ in our analysis:

- Estimating $N\tau$ is confounded with mutation rate $\mu$. Without independent estimates of mutation rate, our estimates would be highly uncertain.

- Estimating $N\tau$ from single-timepoint data is easiest when low-VAF variants are available (VAF < $\sim 0.01$), where the expected VAF density distribution converges to $N\tau\mu$. The sequencing depth in UK Biobank is insufficient to detect these variants.

Throughout this analysis, therefore, we simply used $N\tau = 100{,}000$ cell-years. Allowing $N\tau$ to vary across different individuals with a mean of $100{,}000$ is unlikely to affect our inferences. Since $N\tau$ enters the model through the term $N\tau\mu$, overestimation of $N\tau$ would lead to an underestimation of $\mu$, but the effect on estimates of fitness would be negligible. Hence, uncertainty in $N\tau$ (or variation in $N\tau$ among individuals) is unlikely to affect the inferences materially.

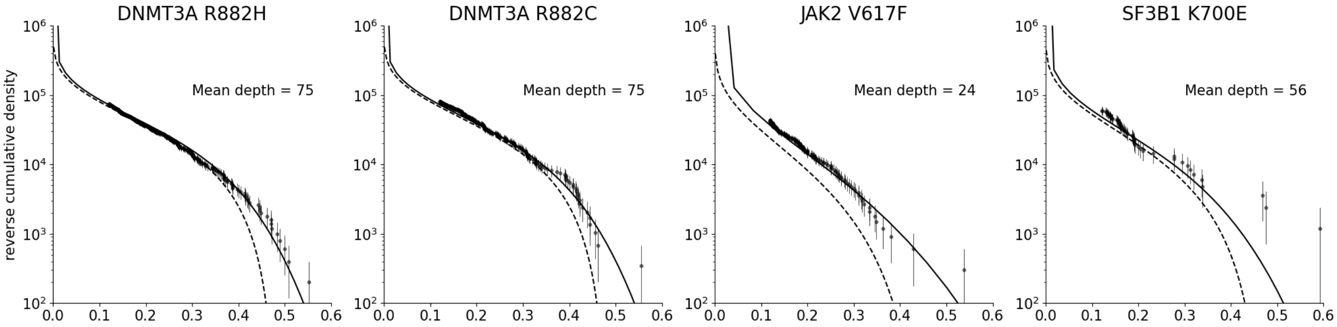


**Fig. S4. Effect of finite sequencing depth on predictions of variant distributions.** Comparison of predictions of variant density using the constant-growth model before accounting for sequencing effects (dashed lines) and after adjusting for finite sequencing depth (solid lines). The adjustment was done using the mean observed sequencing depth for variant carriers (depth shown in each plot). Lower depth results in a more pronounced sequencing adjustment.

**C. Adjusting for sequencing depth.** The estimates of VAF are subject to measurement error due to finite sequencing depth. We adjusted eq. 2 to account for the sequencing depth in UKB by modelling the number of variant reads as a binomial distribution $\text{Bin}(D, f)$, where $f$ is the true VAF and $D$ is the depth. The probability of finding $r$ variant reads is then

$$\rho_d(r|D,t,\{a_i\}) = \int_0^{0.5} P(r|D,f)\ \rho(f,t)df \qquad \textbf{(3)}$$

Where

$$P(r|D,f) = \binom{D}{r} f^r (1-f)^{D-r}$$

The effect of sequencing depth on a selection of CH hotspot variants in UKB (under the constant growth model) is shown in Fig. S4. Notably, accounting for sequencing depth predicts that some variants should appear at $f > 0.5$ due to sampling noise. For each variant, the value of $D$ was taken to be the mean sequencing depth for variants at that position. While this method is theoretically applicable for any sequencing depth, in practice some variants with very low average sequencing depth in UKB (*JAK2* V617F and *GNB1* K57E, average depth $\sim 20$) were excluded since the low depth led to unreliability in parameter estimation.

The distributions of VAF shown in all figures in the main text are *reverse cumulative* distributions, with the density values divided by mutation rate. This format was chosen to remove the need to sort variants into bins and to achieve greater visual separation between age groups. The reverse cumulative density $\mathrm{C}(r|D,t)$ is calculated by summing the discrete distribution $\rho_d(r|D,t)$ in reverse order from $r = D$:

$$\mathrm{C}(r|D,t) = \sum_{r'=D}^{r} \rho_d(r'|D,t)$$

**D. Age-adjusted maximum likelihood parameter estimation.** Together, the age of a variant carrier and the VAF of the variant carry contain joint information about the likely mutation rate and fitness of the variant. Estimating model parameters using a least-squares fit to the overall VAF distribution (Fig. S5a), as in previous studies[10], does not utilise this joint information. We found that least-squares fitting often gives parameter estimates that fit the overall distribution well but the age-stratified distributions poorly (Fig. S14, S15). In order to fit a model across all ages, we estimated the parameters $\{a_i\}$ for each model using a maximum-likelihood approach that explicitly accounted for the age of each individual carrying a variant (Supplementary Fig. S5b). For a single individual $j$ with a CH variant, the likelihood of the parameters $\{a_i\}$ (e.g. for the constant-growth model, $\{a_i\} = \{\mu, s\}$), given the data, is simply the probability of finding the variant at its observed read depth $r_j$, given the age of the participant:

$$\mathcal{L}_j(\{a_i\}|r_j, D_j, t_j) = \rho_d(r_j|D_j, t_j, \{a_i\})$$

There is also a contribution to the likelihood from individuals *without* variants. An individual with no observed variant could either actually have no variant or carry a variant below the detection limit in the UKB data. We had determined this limit $f_0$ for each variant by trimming the data (described in section 1A). The contribution to the likelihood of an individual $k$ without a variant is:

$$\mathcal{L}_k(\{a_i\}|t_k) = \int_0^{f_0} \rho(f|t_k, \{a_i\})df$$

using the continuous VAF distribution for the appropriate model (e.g. eq. 2).

The total log-likelihood is therefore:

$$\log\mathcal{L}_{\text{total}}(\{a_i\}|\text{data}) = \sum_{j \text{ in } V} \log\mathcal{L}_j(\{a_i\}|r_j, D_j, t_j) + \sum_{k \text{ not in } V} \log\mathcal{L}_k(\{a_i\}|t_k) \quad \textbf{(4)}$$

Where $V$ is the set of individuals in whom variants are observed. Parameters $\{a_i\}$ were estimated by minimising this log-likelihood using a Nelder-Mead simplex algorithm[43] implemented in Python.

The maximum likelihood estimation (MLE) approach has a number of advantages over least-squares fitting. Since the likelihood incorporates all aspects of the data (including VAF and age for each individual), MLE implicitly fits VAF distributions to all ages at once. This is demonstrated in Fig. S5c-d: with least-squares fitting (C) the model appears to fit well to the overall density, but when stratified by age the model fits poorly. With maximum-likelihood estimation (D), the model attempts to fit parameters that explain the distribution at all ages, and it is clear both on aggregate and when stratified by age that the constant-growth model is a poor fit for the data. Furthermore, the MLE approach removes the need for arbitrary binning by VAF or by age. The age divisions presented here were chosen solely for visualisation purposes – the model fitting approach is agnostic to this choice.

**E. Uncertainty in parameter estimation.** The uncertainty in the parameter estimates for the maximum-likelihood method was obtained using profile likelihoods. Usually the parameter of interest was the ratio of early-life to late-life growth ('deceleration factor') $\alpha = s_1/s_2$. To obtain the uncertainty in $\alpha$, the likelihood function was redefined as a function of $\mu$, $s_1$, and $\alpha$ (which does not alter the maximum-likelihood estimates). The profile likelihood was then constructed by varying $\alpha$ stepwise away from $\hat{\alpha}$ and finding the likelihood-maximising values of $\mu$ and $s_1$ for each step[44].

Wilks' theorem states that for a likelihood ratio (the difference between two log-likelihoods), the test statistic $\lambda$ is $\chi^2$-distributed[45], where

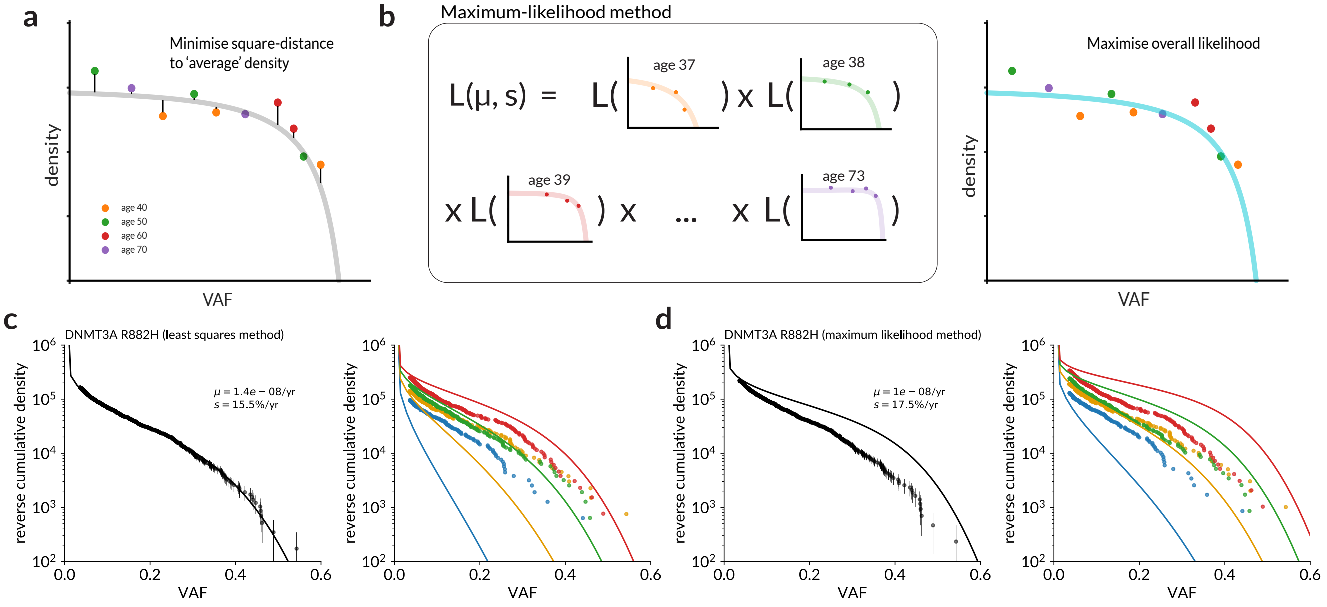


**Fig. S5. Age-adjusted maximum likelihood parameter estimation (A)** Simple least squares fitting estimation ignores the information from the ages of the variant-carrying individuals **(B)** A maximum-likelihood approach, evaluating the likelihood for each individual variant given the (integer) age of the carrier, utilises the joint information from age and VAF. **(C)** Predicted VAF distribution for UKB *DNMT3A* R882H, using a constant growth model fitted to the overall VAF distribution with a least-squares approach (overall (left) and stratified into four age quartiles (right). **(D)** Predicted VAF distribution for UKB *DNMT3A* R882H, using a constant growth model fitted using age-adjusted maximum likelihood (overall (left) and stratified into four age quartiles (right).

$$\lambda = 2\Delta \log \mathcal{L} = \log \mathcal{L}(\hat{\boldsymbol{\theta}}) - \log \mathcal{L}(\boldsymbol{\theta}) \quad \textbf{(5)}$$

Where $\boldsymbol{\theta}$ represents the model parameters, and $\hat{\boldsymbol{\theta}}$ represents the values of the model parameters that maximise the likelihood. With one degree of freedom, the 95% confidence level of the $\chi^2$ distribution is $\lambda = 3.84$ (Fig. S6).

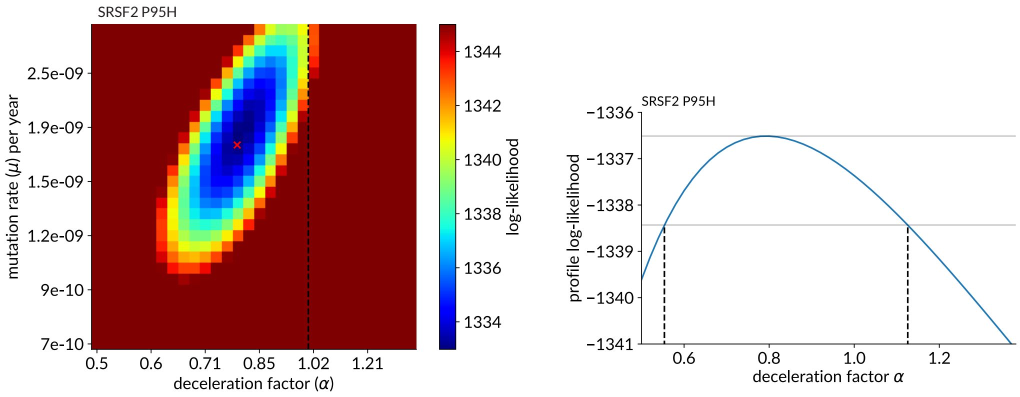


**Fig. S6. Confidence interval estimation using profile likelihood. Top:** Two-dimensional cross-section of log-likelihood function $\mathcal{L}(\mu, s_1, \alpha)$ for the two-phase CH model for SRSF2 P95H. $s_1$ dimension not shown. Position of maximum-likelihood estimate marked with red 'x'. **Bottom:** Profile log-likelihood for deceleration parameter $\alpha$, obtained by varying $\alpha$ and re-maximising the likelihood allowing only $\mu, s_1$ to vary. 95% confidence interval in $\alpha$ (dashed lines) is defined where this function falls to 1.92 below the maximum (solid grey lines).

**F. Developmental mutations.** Earlier The above modelling describes a mature, static haematopoietic system with a steady HSC population ($N\tau = 100{,}000$ cell-years). It is also possible to acquire somatic clones in development (while the HSC population is growing) – if sufficiently prevalent, developmental clones could contribute to a higher 'baseline prevalence' of CH in young people and give the appearance of deceleration with age. Poon et al.[14] calculated the density distribution of these clones (for a birth-only process without logistic correction) as:

$$\rho(f) = \frac{\mu_d}{\log 2} \frac{1}{2f^2} \quad \textbf{(6)}$$

where $\mu_d$ is the developmental mutation rate (mutations per cell doubling). This is estimated in the same publication to be no greater than 2-4 mutations per cell doubling across the whole genome.

***Developmental contribution at individual hotspots.*** A mutation rate of 2-4 per cell doubling across the genome corresponds to $\sim 10^{-9}$ per cell doubling at a single position. Considering that the real density falls to zero as $f \to 0.5$, the reverse cumulative density at the VAF detection limit $f^*$ is approximately

$$\frac{\mu_d}{\log 2}\frac{1}{2f^*}$$

For $f^* = 0.1$, this corresponds to a density of the order $1 - 10$ (after normalising by total mutation rate as in the plots presented here). Even allowing for some variation in developmental mutation rate, this value is several orders of magnitude lower than the variant densities observed in UKB, and suggests that developmental mutations are unlikely to play a major role in the age-related deceleration at hotspots.

***Developmental contribution across whole genes.*** At the gene-aggregated level, if developmental mutations contributed significantly to apparent CH in a gene this would produce 'static' age behaviour similar to that observed in genes such as KMT2C and NF1, among others (Supp. Note 1B.2). Longer genes would be expected to accumulate more developmental mutations, so we calculated the 'effective coding length' of each gene by applying a subset of the exome-wide filters (Supp. Note 1B) to every combination of possible SNVs along the coding length of each gene, identifying where a hypothetical variant could be called as a putative CH driver (Fig. S7). Some artifact filters could not be applied to hypothetical variants (such as the strand bias or population-frequency filters), so this estimate is conservative (i.e. likely to overestimate the true 'effective coding length'). We compared the observed VAF distribution for each gene, normalised by effective coding length, with the expected distribution for developmental mutations based on Eq. 6 (Fig. S8). The observed VAF densities were at least an order of magnitude larger than would be expected from developmental mutations at a rate of 2-4 mutations per cell doubling. Although this is an approximate analysis (not considering, for instance, variability in the mutation rate between different sites), the size of the discrepancy suggests that the mutations observed in UKB that do not exhibit clonal growth are not developmental in origin, but are more likely to be artefactual. The estimates for both the mutation rate (2-4 per cell doubling) and the effective gene lengths are more likely than to be overestimates than underestimates, so the real discrepancy is likely to be even larger than seen here.

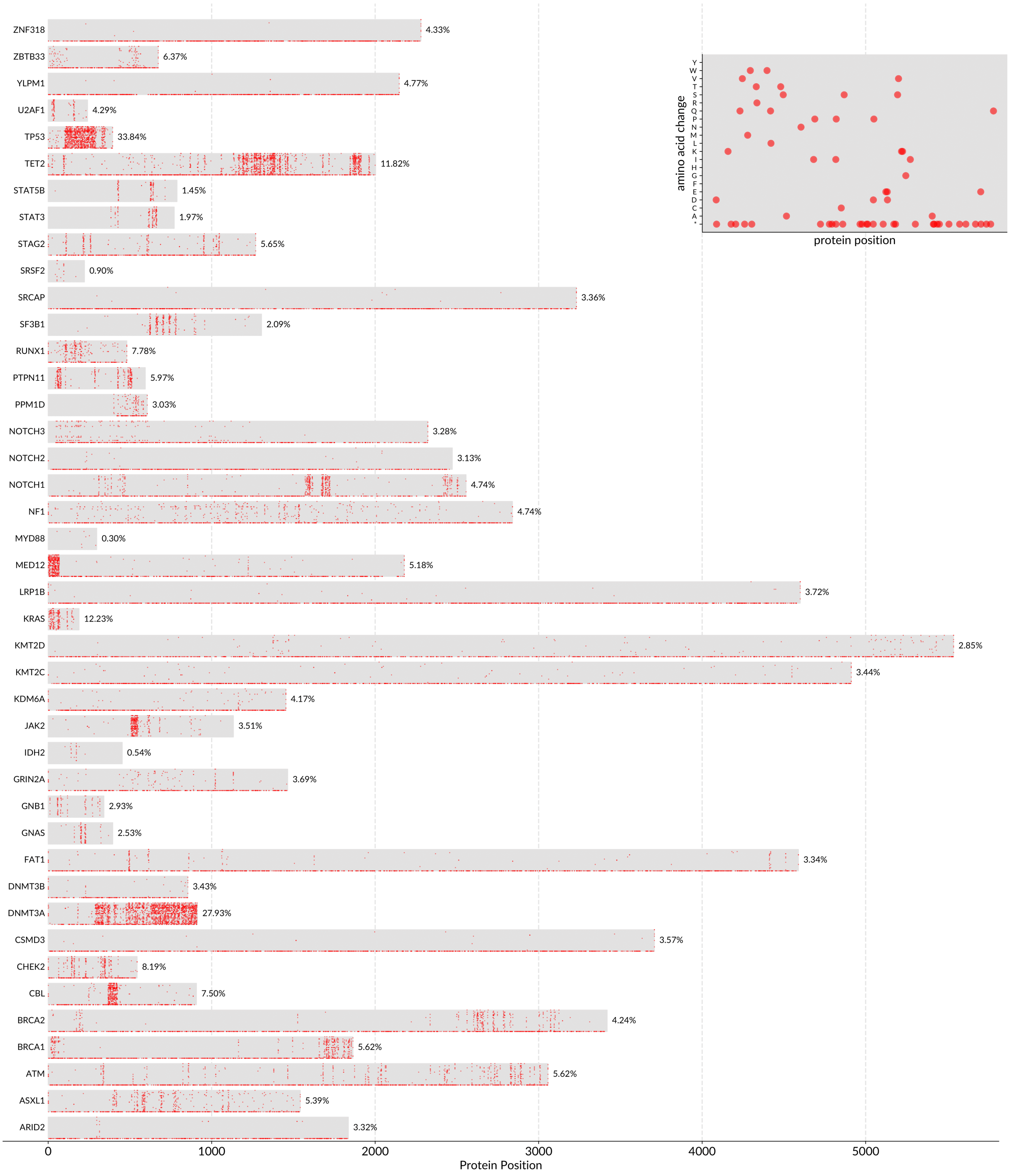


**Fig. S7. Estimation of effective CH coding length.** Grey shaded area shows coding length of each gene (amino acids in the canonical transcript), red dots mark where a hypothetical variant, if observed, could qualify as a 'putative driver' based on the position-based filters outlined above. Vertical positions of red dots represent specific amino acid substitutions (as shown in **inset** with example data). Asterisk represents a stop codon. The fraction of all possible single nucleotide substitutions that could pass the position-based filters is shown to the right.

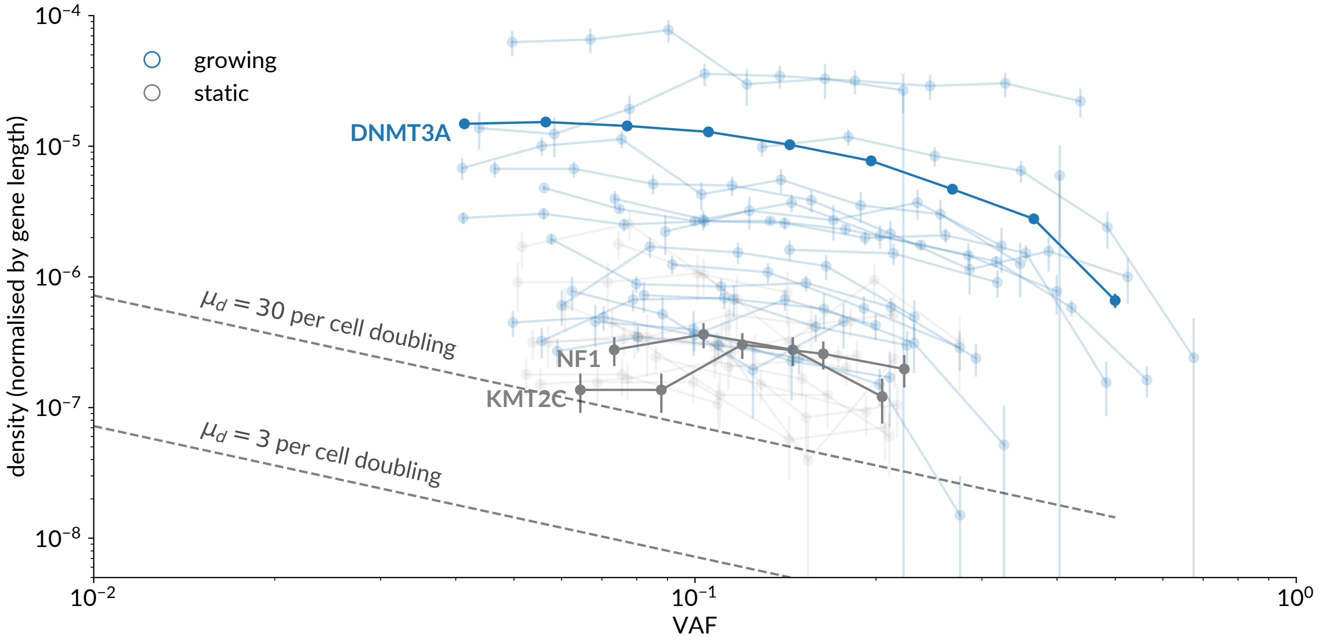


**Fig. S8. Evaluating the contribution of developmental mutations** VAF distributions (non-cumulative) for top 40 genes observed in the exome-wide CH calling data, normalised by 'effective gene length'. Genes identified as 'growing' (positively associated with age with $p < 0.05$) are coloured blue, others are coloured grey. Dashed lines show expected contribution from developmental mutations for a mutation rate of 3 per cell doubling (approximately the estimated rate by Poon et al.[14]) and tenfold higher.

### Supplementary Note 3: Age-dependent models of clonal dynamics

This section contains the details of three mathematical models used in this analysis, to generalise the constant growth model:

A. **Two-phase model of clonal dynamics:** To quantify the decelerating behaviour of different CH drivers, a model assuming different growth rates for early ($s_1$) and later life ($s_2$).

B. **Variation in fitness between individuals:** To assess the inter-individual variability in fitness, a model in which the fitness effect conferred by a CH hotspot variant is variable across the population.

C. **Clonal competition:** To assess the potential role of clonal competition in age-dependent CH dynamics, a model in which clonal expansion is constrained by the expansion of other clones experiencing positive selection in the same individual.

In each case, we started from the constant-growth model (eq. 2) and derived a new density distribution (and corresponding likelihood function) with alternative parameters $\{a_i\}$.

**A. Two-phase model of clonal dynamics.** We developed a two-phase model of clonal dynamics in order to quantify the deceleration (or acceleration) of clones with age. The youngest individual in the UKB cohort is 37, and there are only 3 individuals below the age of 40. As a result, inferences about clonal dynamics before age 40 are limited to estimating an average growth rate up to this point. We therefore modelled stem cell ageing as a two-stage process with a constant fitness effect $s_1$ until $t = 40$ yrs and a second fitness effect $s_2$ thereafter. We defined a 'deceleration factor' $\alpha = s_1/s_2$ representing the change in fitness – a larger deceleration factor represents a more pronounced deceleration.

The fitness parameter $s$ enters the model (eq. 2) through the characteristic clone size $\tilde{n}(t)$. Therefore, we can obtain an expression for the variant density in a scenario with variable clonal fitness by adapting $\tilde{n}(t)$ to account for the history of the growth rate (similarly to Watson et al.[10], Supplementary Material 3):

$$\tilde{n}(t) = \frac{\exp(st) - 1}{s} \quad \rightarrow \quad \frac{\exp(\int_0^t s(t')dt') - 1}{s(t=0)}$$

which simplifies to the constant-growth expression for constant $s$. The expression for variant density is then simply eq. 2 with $\tilde{n}$ substituted as above. We validated this approach with simulations (Fig. S9a-b). The resulting expression has one additional free parameter: $\{\mu, s\} \rightarrow \{\mu, s_1, s_2\}$. We fitted the model using the maximum-likelihood method already described, estimating the maximum-likelihood values of the three parameters independently for each hotspot variant (in the hotspot analysis) or for each gene (in the exome-wide analysis) (Fig. S16).

***Interpretation of the two-phase model.*** In this model, the late-life fitness effect $s_2$ can be interpreted as an average clonal expansion rate over the age range of the UKB participants (approximately 40-70). The interpretation of the early-life fitness effect $s_1$ is more subtle; since the youngest participants in the UK Biobank are around 40 years of age, it was not possible to determine precisely how clonal fitness changes with age: for example, the fitness could change abruptly in childhood or early adulthood, or decline steadily. In the two-phase model outlined above, $s_1$ is the growth rate before age 40 years, and provides a lower bound on the growth rate in early life before age 40. If the deceleration happens earlier than age 40, then the early-life growth rate would need to be greater than $s_1$ to produce the same VAF distributions over the observed period.

A constant growth rate followed by a step change at age 40 is very unlikely to be true in practice. However, since the parameter $\tilde{n}(t)$ depends primarily on the time-integral of $s$, changing the fitness profile before age 40 has only a modest effect on the observed VAF densities as long as $\frac{1}{s_0}\int_0^{40} s\,dt$ remains constant ($s_0$ is the fitness at $t = 0$). Where differences arise, they are due to the factor of $s$ in the denominator (which relates to the transition from the early stochastic growth phase to 'established' clonal growth). Thus, more noticeable changes to the distributions occur in scenarios where, for example, growth is all concentrated in a very short period in early life.

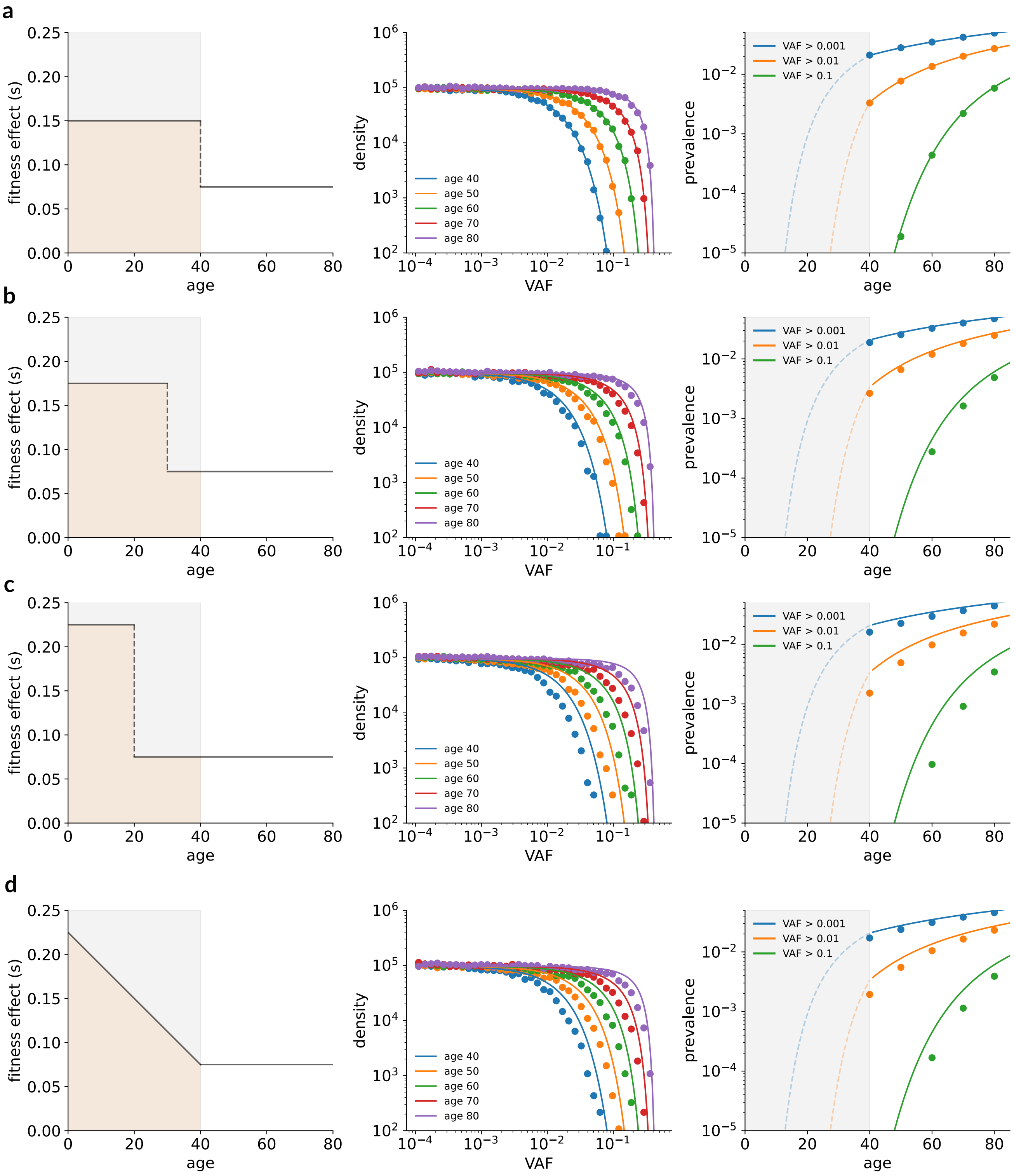


**Fig. S9. Two-stage models of clonal dynamics.** Comparison of simulated data (points) and theoretical expectations (lines) using a two-stage model of clonal dynamics, and comparing the effects of different fitness profiles before age 40 on observable variants. **Left:** Examples of different possible fitness profiles before age 40. **(a)** shows the profile used in the main analysis. The beige shaded area is held constant across the different profiles. **Middle:** VAF densities derived from agent-based simulations of clonal dynamics for each fitness profile (points) compared with theoretical expectation *from the topmost profile (a)* (lines), showing the modest effect of different fitness profiles before age 40. **Right:** prevalence of variants derived from simulations at different limits of detection (points) compared with theoretical expectations from the topmost profile (lines)

### B. Variation in fitness between individuals.

***B.1. Apparent deceleration from a distribution of fitness effects.*** This model is motivated by the idea that across a population of individuals carrying the same variant clone, the fitness effect of that clone may vary. We modelled this scenario by drawing the fitness effect $s$ for each individual from a distribution of fitness effects (DFE) $g(s)$. To see how this could give rise to age-related deceleration, it is instructive to consider a simple example: a DFE consisting of two delta functions at $s = 0.14$ and $s = 0.18$, with the higher-fitness variant ten times rarer (Fig. S10a):

$$g(s) \propto \delta(s-0.14) + \frac{1}{10}\delta(s-0.18)$$

The expected VAF distribution for this DFE is the weighted sum of the two VAF distributions for each delta function, obtained using eq. 2 (since a delta-function DFE is equivalent to constant exponential growth). The resulting VAFs are shown in Fig. S10b – at younger ages, the contribution from the higher-fitness variant dominates (as the lower-fitness clones have not had time to grow yet), but as age increases, the lower-fitness clones increasingly contribute to the observed distribution. If observed at a particular VAF window, the result can appear similar to a deceleration as the distribution becomes increasingly dominated by slower-growing variants. The VAF distribution also changes shape, with the characteristic fall-off at $n \approx \tilde{n}$ becoming more gradual due to the presence of different values of $\tilde{n}$ within the population.

To evaluate whether variation in fitness could explain the UK Biobank data, we fitted a more plausible model with a continuous distribution of fitness effects. We assumed fitness was distributed normally about a mean fitness $\bar{s}$ with standard deviation $\sigma$. This choice of distribution was motivated by the principle that a given CH driver variant should confer some 'intrinsic' fitness to an HSC, but that fitness may be modified by a combination of other factors (such as germline genetic effects) specific to each individual. The overall density is obtained by integrating the constant-growth density (eq. 2) over the normal distribution:

$$\rho(f,t,\{\mu,\bar{s},\sigma\}) \propto \int_0^\infty \rho(f,t,\{\mu,s\}) \, \exp\left[-\frac{(s-\bar{s})^2}{2\sigma^2}\right] ds \tag{7}$$

As $\sigma \to 0$, the density tends towards eq. 2 as expected (Fig. S10d). In practice, the numerical integration to infinity was unstable (especially for small values of $\sigma$, so integration limits were modified to $\bar{s} \pm 5\sigma$ (note that negative $s$ is not inherently problematic: clones with a negative fitness simply go extinct over the long term so have no effect on observable VAF distributions). We validated the theory with simulations (Fig. S10f). Comparing Fig. S10d and S10e demonstrates the apparent deceleration caused by a Gaussian distribution of fitness effects. We fitted eq. 7 to the UK Biobank data, inferring $\bar{s}$ and $\sigma$ independently for each decelerating CH hotspot variant using the maximum-likelihood approach described above.

***B.2. Detection limit for fitness.*** The lower detection limit for VAF was found to be around 3-4% VAF for most hotspots. This imposes a natural lower limit of detection for $s$: if a variant does not confer sufficient fitness to achieve the minimum VAF, it will never be detected. This limit decreases with age as clone have more time to grow. Since the largest variants have VAF $f \sim \tilde{n}(t)/2N$, the lowest detectable fitness at age 40 is around $s = 17\%$ per year, falling to around $9\%$ per year at age 70. Consequentially, this impedes our ability to characterise a distribution of fitness effects: since we cannot see the effects of any variants below the limit, we cannot make assumptions about the DFE below the limit. This can be illustrated using the the DNMT3A hotspots: the best-fit DFEs imply an implausibly large single-site mutation rate to low-fitness variants[10] (Fig. S11a, red), but another DFE can be found (green) that produces very similar VAF distributions (Fig. S11b-c) with much more plausible parameters. Inspecting the two distributions, it is clear that only higher-$s$ portion matters, and the large number of low-fitness variants have negligible effect as they cannot be detected in UKB. This illustrates the limitations of using low-resolution sequencing data to infer distributions of fitness effects.

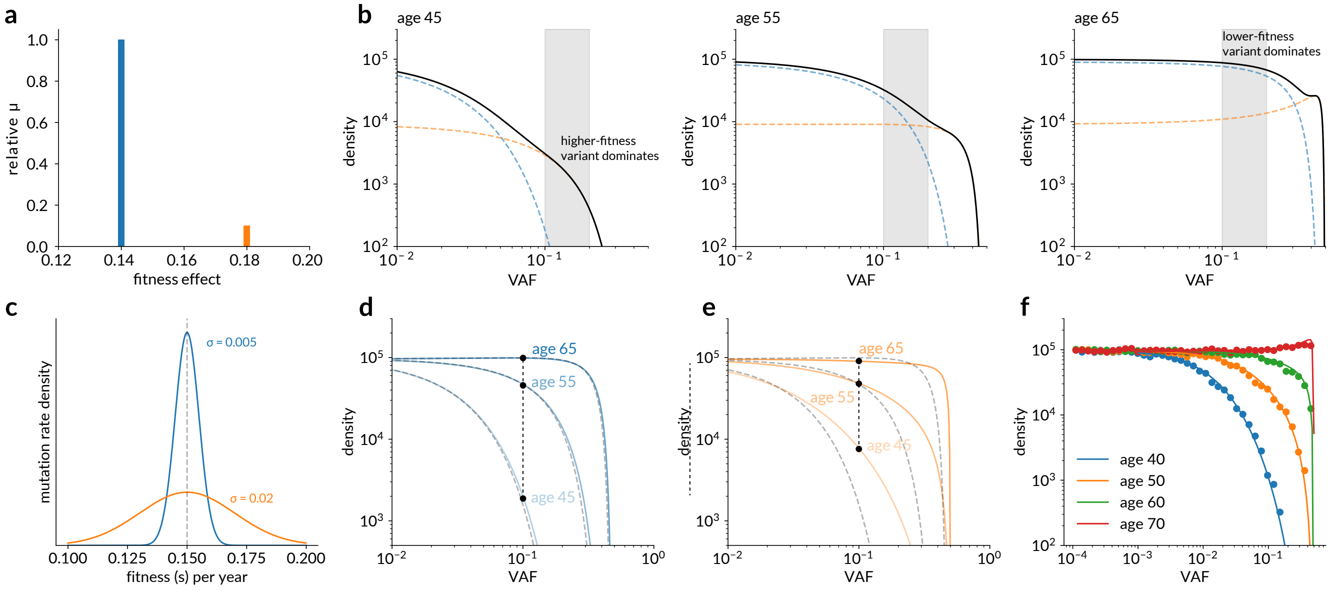


**Fig. S10. Distribution of fitness effect influences VAF distribution (a)** Example DFE constructed from two delta functions at $s = 0.14$ and $s = 0.18$. **(b)** Predictions of VAF density distribution for the two-delta-function DFE, showing the contribution of each delta function (dashed lines) to the overall distribution (black line) at ages 45, 55, and 65. Examining the shaded VAF range demonstrates how the distribution is dominated by high-fitness clones at younger ages, and by lower-fitness clones at older ages. **(c)** Example DFEs of different widths: $\sigma = 0.005$ (blue) and $\sigma = 0.02$ (orange). Dashed line shows mean fitness $\bar{s} = 0.15$ per year. **(d-e)** VAF density at ages 45, 55 and 65 for the two DFEs shown in (c), each compared to a density for a constant-fitness model with $s = 0.15$ (dashed lines). A broad DFE (e) can produce VAF distributions that resemble deceleration, as shown by comparing the density at different ages at VAF = 0.1 (black markers) **(f)** Simulations (points) of a DFE model with $\mu = 1 \times 10^{-7}$, $\bar{s} = 0.15$, $\sigma = 0.02$ compared with theoretical expectation (lines) for a range of ages.

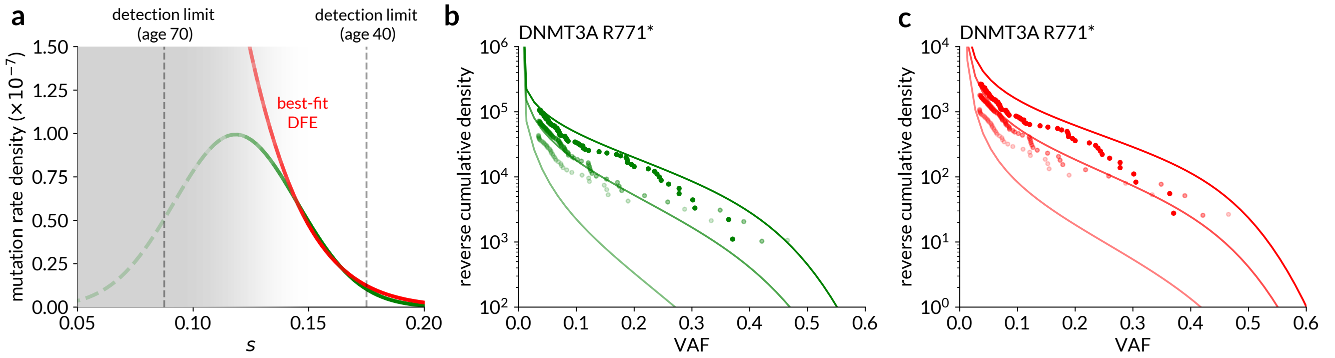


**Fig. S11. Limits of detection for distribution of fitness effects. (a)** Two example DFEs for *DNMT3A* R771*: the best-fit DFE (red) and another DFE with similar characteristics at high $s$ but more plausible parameters (green). Limits of detection for $s$ are shown for age 40 and 70 (dashed lines) and the decreasing contribution of the low-$s$ portion of the DFE is indicated visually with shading. **(b-c)** Reverse cumulative VAF distributions for *DNMT3A* R771* compared with predictions from the green **(b)** and red **(c)** DFEs. The VAF distributions are similar despite the substantially different DFEs.

**C. Clonal competition.** Clonal expansion is constrained by the expansion of other clones experiencing positive selection in the same individual. This is a diverse scenario that could encompass competition between numerous clones driven by mutations, copy number events or other unknown drivers of clonal expansion. We considered a simple model to assess whether it was plausible that pervasive clonal competition gives rise to clonal deceleration in early life. In this model, an observable variant (e.g. *DNMT3A* R882H) with parameters $\{\mu_1, s_1\}$ competes with a generic 'background', which we treat as a single event with a mutation rate $\mu_2$ and fitness effect $s_2$. The background need not be observed. The model accounts for the possibility of multiple background clones arising independently in the same individual, but we assume that these events cannot occur in the same clone (i.e. no double-mutant clones). The model, while very simple, is useful for its mathematical tractability - we demonstrate further below that the conclusions we can draw from the single-fitness background model still hold for more complex models of competition (e.g. with a distribution of fitness in the hidden drivers).

In the simple model expression for the expected variant density is obtained by calculating the frequency of the observed clone in the presence of the expanding background clone. The expected distribution of total observed clone size $n_1$ in a population *without* competition is given by eq. 1, repeated here:

$$\rho_n(n_1|t,\{\mu,s\})dn = n_1^{\theta-1}\frac{\exp(-n_1/\tilde{n_1}(t))}{\Gamma(\theta)\tilde{n_1}(t)^{\theta}}dn_1$$

with $\theta$ and $\tilde{n}(t)$ as before. Considering an HSC population that started with $N$ wild-type cells, with an additional observable clone of size $n_1$ and background clone of size $n_2$, the observable VAF $f_1$ is defined as

$$2f_1 = \frac{n_1}{N+n_1+n_2}$$

Changing variables in eq. 1 from $n_1$ to $f_1$ gives

$$\rho_f(f_1)df = \frac{(2(N+n_2))^{\theta}}{f^{1-\theta}(1-2f)^{1+\theta}}\exp\left(\frac{-2(N+n_2)f}{\tilde{n}(t)(1-2f)}\right)\frac{1}{\Gamma(\theta)\tilde{n}(t)^{\theta}}df \tag{8}$$

The expected density of observable clones is obtained by integrating eq. 8 over the expected density of background clone sizes $n_2$, given again by eq. 1 with parameters $\{\mu_2, s_2\}$:

$$\rho(f_1|t,\{\mu_1,s_1,\mu_2,s_2\})df_1 = \int_0^{\infty}\rho_f(f_1,n_2)\,\rho_n(n_2)\,dn_2\,df_1 \tag{9}$$

Integrating eq. 9 to infinity was found to be numerically unstable, so the upper limit was chosen as $50\tilde{n}(t)$ in practice, a size far larger than any background clone would be expected to reach. This model has two additional parameters compared with the constant growth model: the background mutation rate and fitness $\{\mu_2, s_2\}$. We validated eq. 9 with simulations (Fig. S12).

For the hotspots and genes which exhibited deceleration, the total log-likelihood was maximised independently in each case using the Nelder-Mead simplex algorithm, estimating $\{\mu_1, s_1\}$ for each variant/gene along with for the $\{\mu_2, s_2\}$ for the background in each case. In a few cases, the model failed to find a maximum (*DNMT3A* Y735C, R729W and R882S, *ZBTB33*, *ATM*).

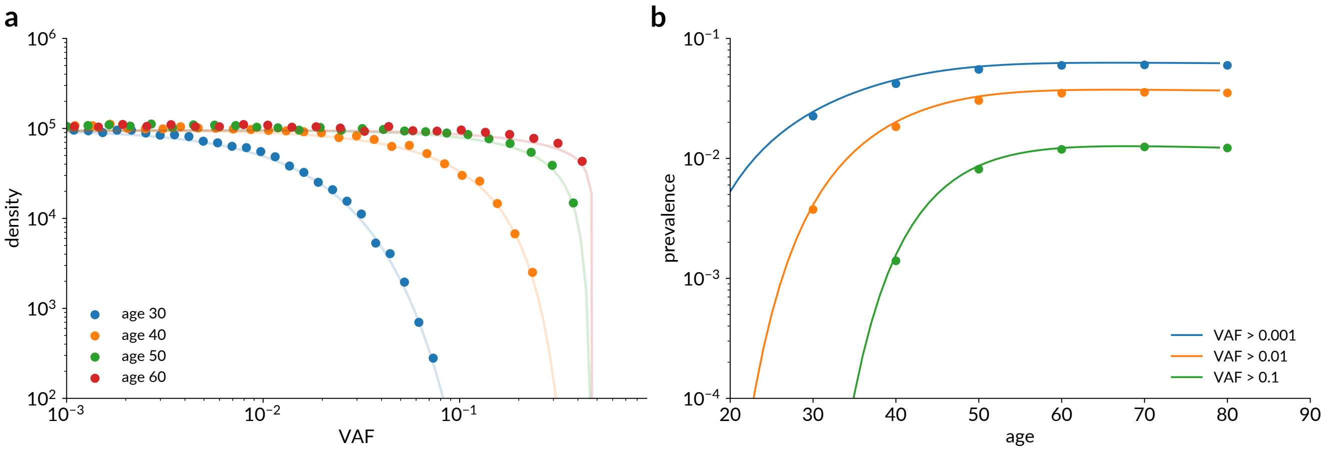


**Fig. S12. Simulations of clonal competition. (a)** Simulated VAF distributions for clonal competition model (points) compared with analytical expectations (lines), for a population identical in size to the UK Biobank cohort (n=416,171) sampled every 10 years from age 30 to 60). Mutation rate, fitness similar to those inferred for DNMT3A R882H. **(b)** Simulated overall prevalence of CH under the clonal competition model (points) compared with analytical prediction (lines), using the same parameters as (a). The lower VAF threshold was varied: VAF > 0.001 (blue); VAF > 0.01 (orange); VAF > 0.1 (green)

***Conditions for clonal competition in the UK Biobank.*** The deceleration of clonal dynamics observed in UK Biobank appears to occur before age 40. For clonal competition to affect clonal dynamics by this age, clones must have sufficient fitness to have expanded to a reasonably large size in this time. The approximate time $t$ between an initial driver mutation occurring and the resulting clone achieving a VAF $f$ is given by

$$\frac{2f}{1-2f} = \frac{n(t)}{N} \approx \frac{e^{st}-1}{N\tau s}$$
$$t \approx \frac{1}{s}\log(1+\frac{2fN\tau s}{1-2f})$$

Using this expression, the minimum fitness $s$ required to achieve VAF $f \sim 0.1$ by age $t = 40$ years is approximately 20-25% per year (assuming $N\tau = 10^5$ cell-years).

For clonal competition to be widespread across the population, the average individual needs to carry at least $\sim 1$ expanded clone. Clones arise at a rate $N\tau s\mu$ per year (since $s\tau$ is the probability of establishment for an individual clone). Therefore, for each individual to have at least an average of 1 mutant clone arise in the first $t_{\mathrm{mut}}$ years of life that survives into middle age, then $N\tau s\mu t_{\mathrm{mut}} \gtrsim 1$. The clone must occur reasonably early in life to have time to grow to large VAF by age 40 (i.e. $st_{\mathrm{mut}} \sim 1$, n), so this requires that $\mu \sim 10^{-5}$ per year or greater. Although these bounds for background fitness adn mutation rate are relatively approximate, they agree with the estimates derived from fitting evolutionary models to the UK Biobank data.

***Extended models of clonal competition.*** Simulations of clonal competition validate the theoretical approach described above. For an observable driver with comparable parameters to *DNMT3A* R882H, competition only becomes important as the mutation rate to the background approaches $1 \times 10^{-5}$ per year (Fig. S13a-c). The simulations confirm that at high mutation rate ($N\tau\mu \sim 1$), we expect multiple clones to contribute towards the unseen background. The effect of this is to relax the constraints on fitness and mutation rate slightly, since early-arising clones are bolstered by smaller contributions from later-arising clones.

Simulations also show that a background consisting of even larger mutation rate to low-fitness clones ($< 10\%$ per year), as suggested by Poon et al.[14], is not sufficient to bring about pervasive clonal competition by middle age (Fig. S13d-e). In this case, despite the large number of driver mutations acquired they do not generally grow to a substantial total VAF nor do they constrain the growth of an acquired *DNMT3A* R882H-like observable driver. This suggests that our conclusions are robust to alternative models of clonal competition. While we have not accounted for the effect of potentially fitness-enhancing 'second-hit' mutations within a clone, Poon et al. estimated that for their estimated distribution the expected number of double mutant clones reaching VAF > 2% by age 60 was less than $10^{-4}$ per person, and so can safely be neglected.

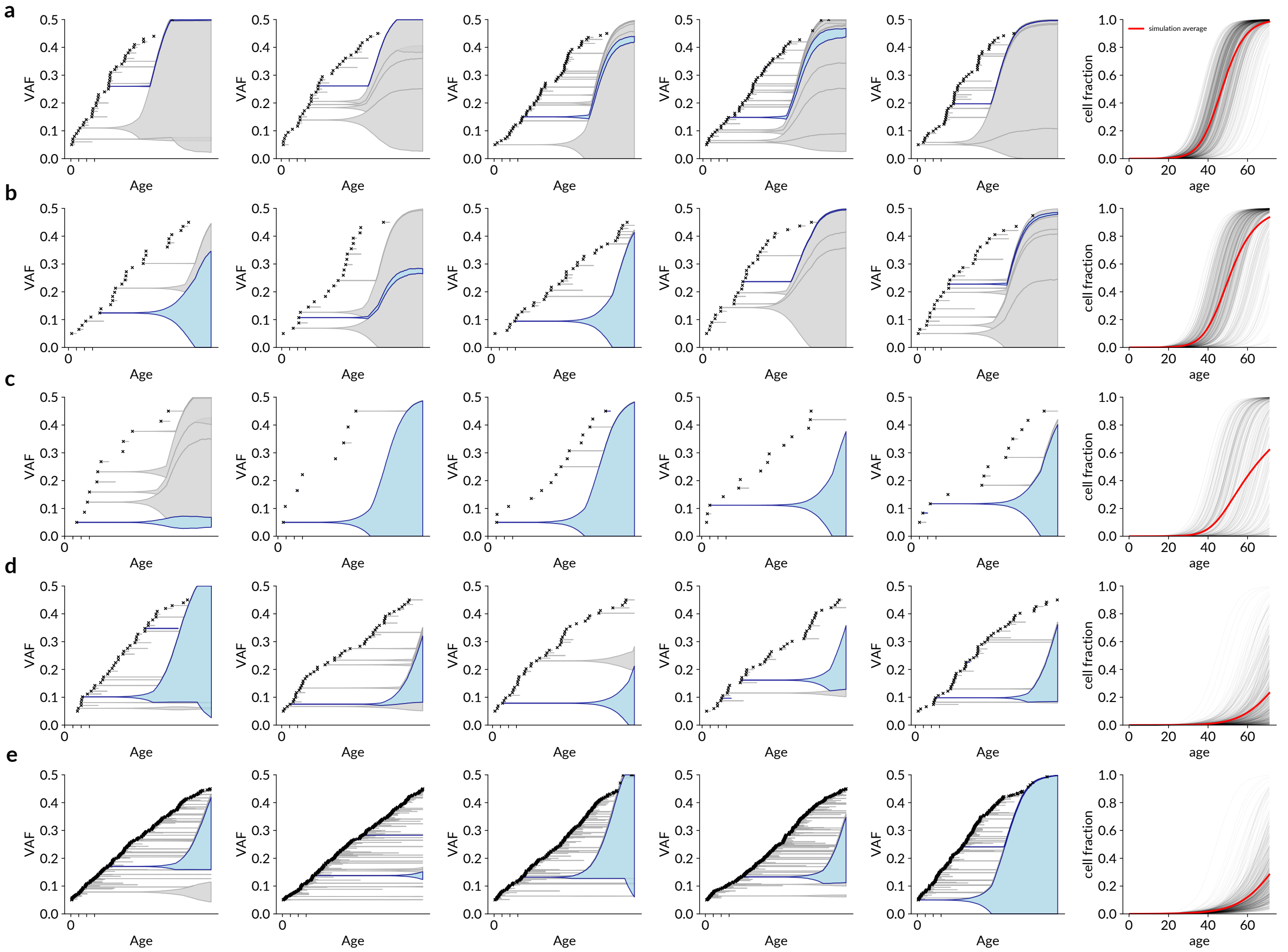


**Fig. S13. Simulations of clonal competition with varying background parameters.** Parameters $(\mu, s)$ for observed drivers (blue) always $(1 \times 10^{-7}, 20\%)$ per year. Simulations conditioned on having at least one long-lived observable clone (blue). Rightmost plot shows proportion of simulated HSCs containing at least one driver (hidden or observable). Grey lines represent 1000 individual simulations, red line represents average across simulations. **(a)** Background $\mu = 1 \times 10^{-5}$ and delta-function fitness $23\%$. **(b)** Background $\mu = 0.6 \times 10^{-5}$ and delta-function fitness $23\%$. **(c)** Background $\mu = 0.2 \times 10^{-5}$ and delta-function fitness $23\%$. **(d)** Background mutation rate and fitness distributed according to stretched exponential distribution described by Poon et al.[14], $p = 3$, $q = 0.1$, with minimum background fitness cut off at $10.5\%$. **(e)** As in (d), but also including more lower-fitness clones (down to a minimum fitness of $3.5\%$)

### Supplementary Note 4: Full results

- Constant-growth model (least squares optimisation): Figures S14 and S15
- Two-stage model: Fig. S16 (hotspots) and Fig. S17 (genes and additional hotspots)
- Distribution of fitness effects model: Fig. S18
- Clonal competition model: Fig. S19 (hotspots) and S20 (genes).

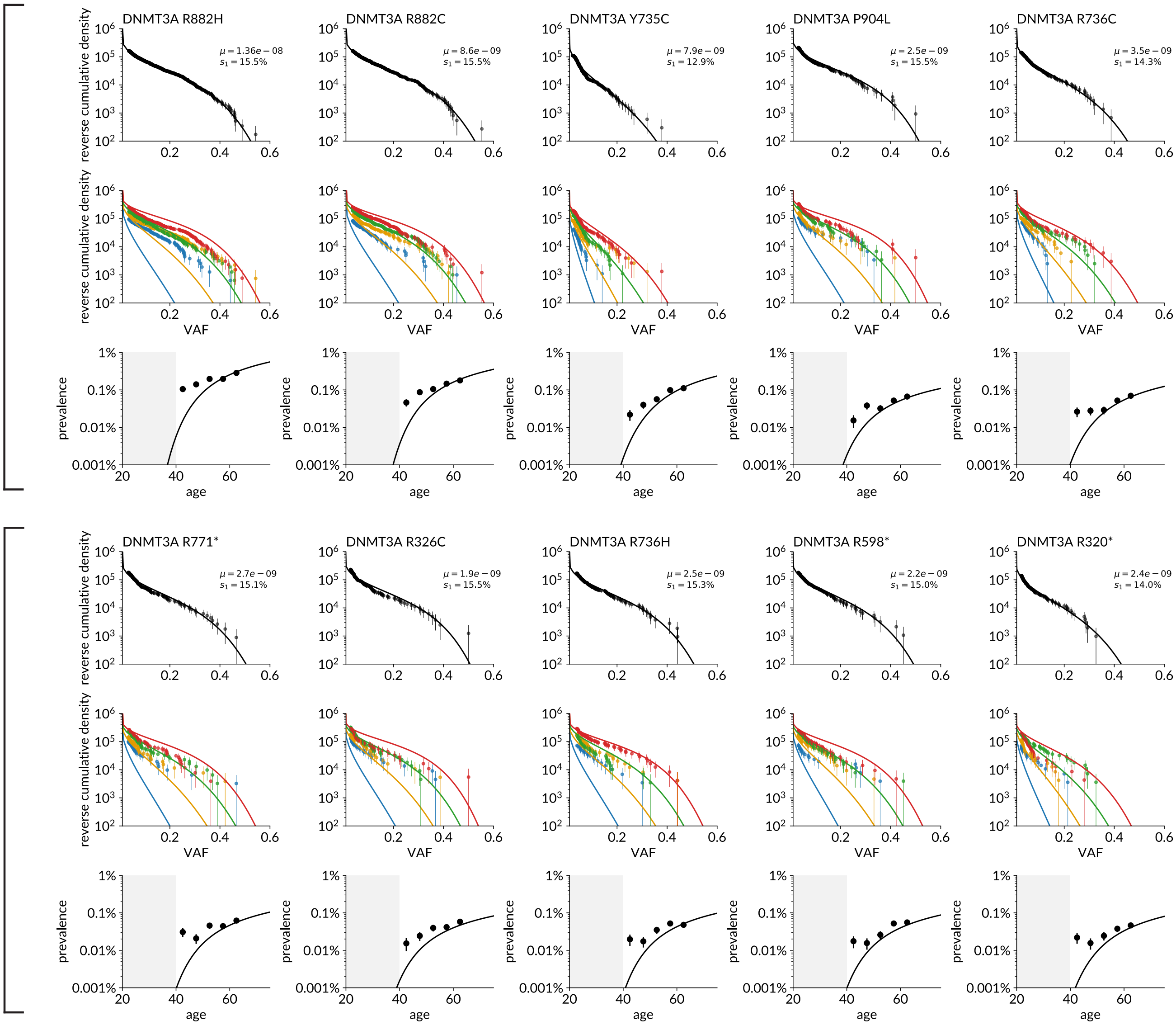


**Fig. S14. Results of constant-growth model fitted to 20 hotspots using least-squares minimisation. (1-10) Top rows in each group:** Reverse cumulative VAF density for each variant, aggregated across all ages (points) compared with predictions from constant-growth model. **Middle rows:** As above, stratified into four age groups. **Bottom rows:** Prevalence of each CH variant by age, compared with predicted prevalence. Grey shaded area shows unobserved period before age 40.

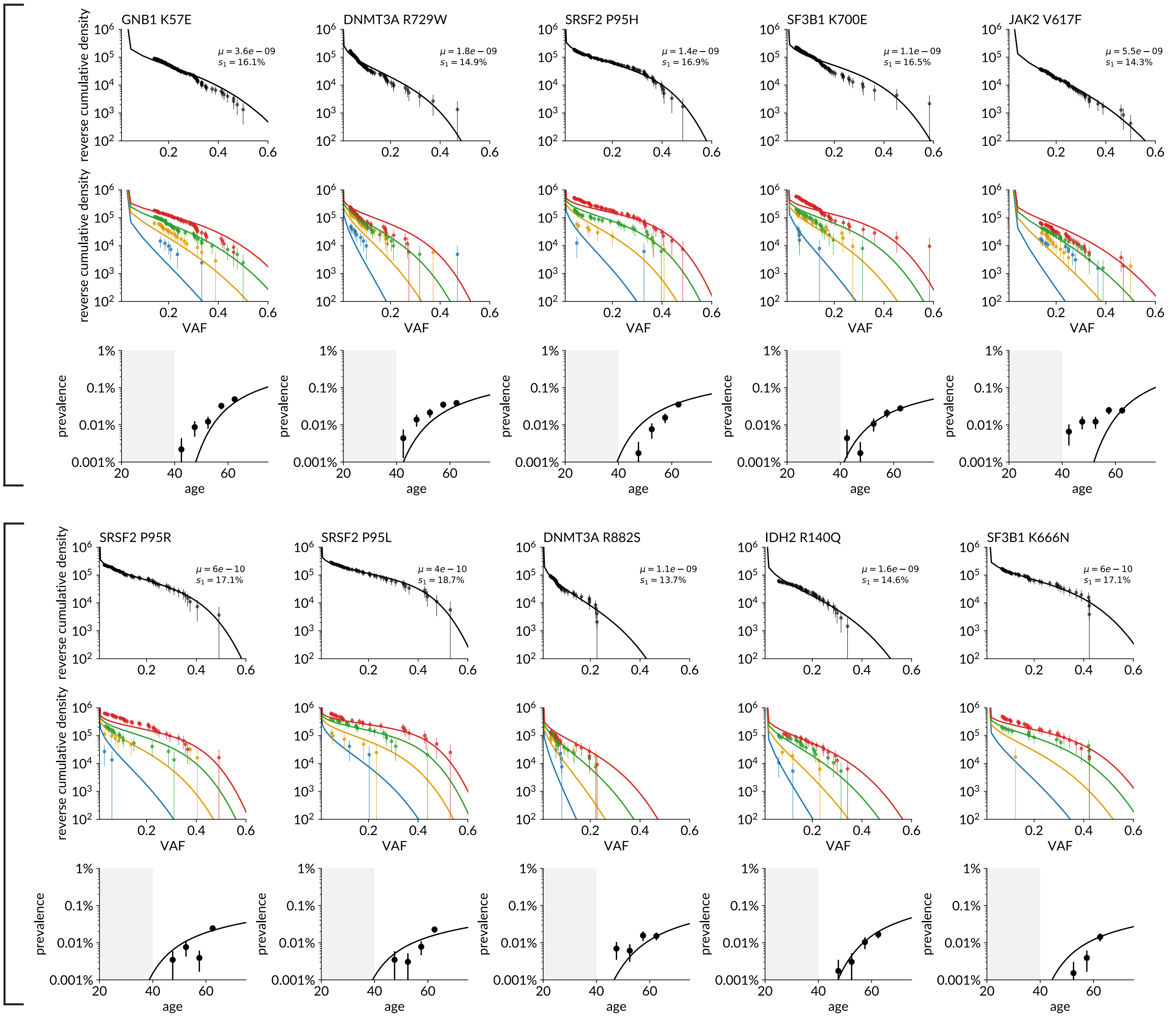


**Fig. S15. Results of constant-growth model fitted to 20 hotspots using least-squares minimisation. (11-20) Top rows in each group:** Reverse cumulative VAF density for each variant, aggregated across all ages (points) compared with predictions from constant-growth model. **Middle rows:** As above, stratified into four age groups. **Bottom rows:** Prevalence of each CH variant by age, compared with predicted prevalence. Grey shaded area shows unobserved period before age 40.

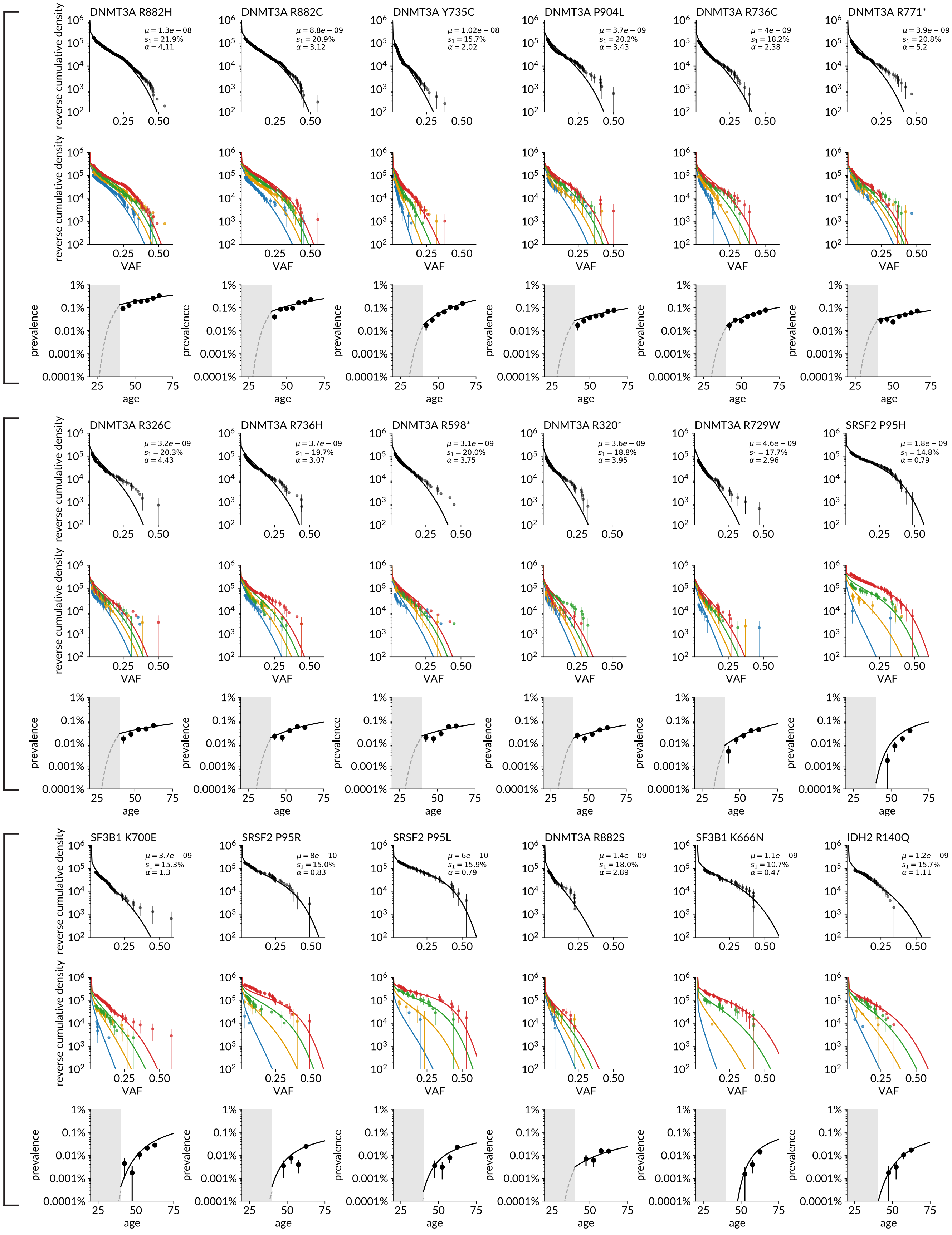


**Fig. S16. Results of two-stage model applied to 18 CH hotspots. Top rows in each group:** Reverse cumulative VAF density for each variant, aggregated across all ages (points) compared with predictions from two-stage model. **Middle rows:** As above, stratified into four age groups. **Bottom rows:** Prevalence of each CH variant by age, compared with predicted prevalence from two-stage model. Grey shaded area shows unobserved period before age 40.

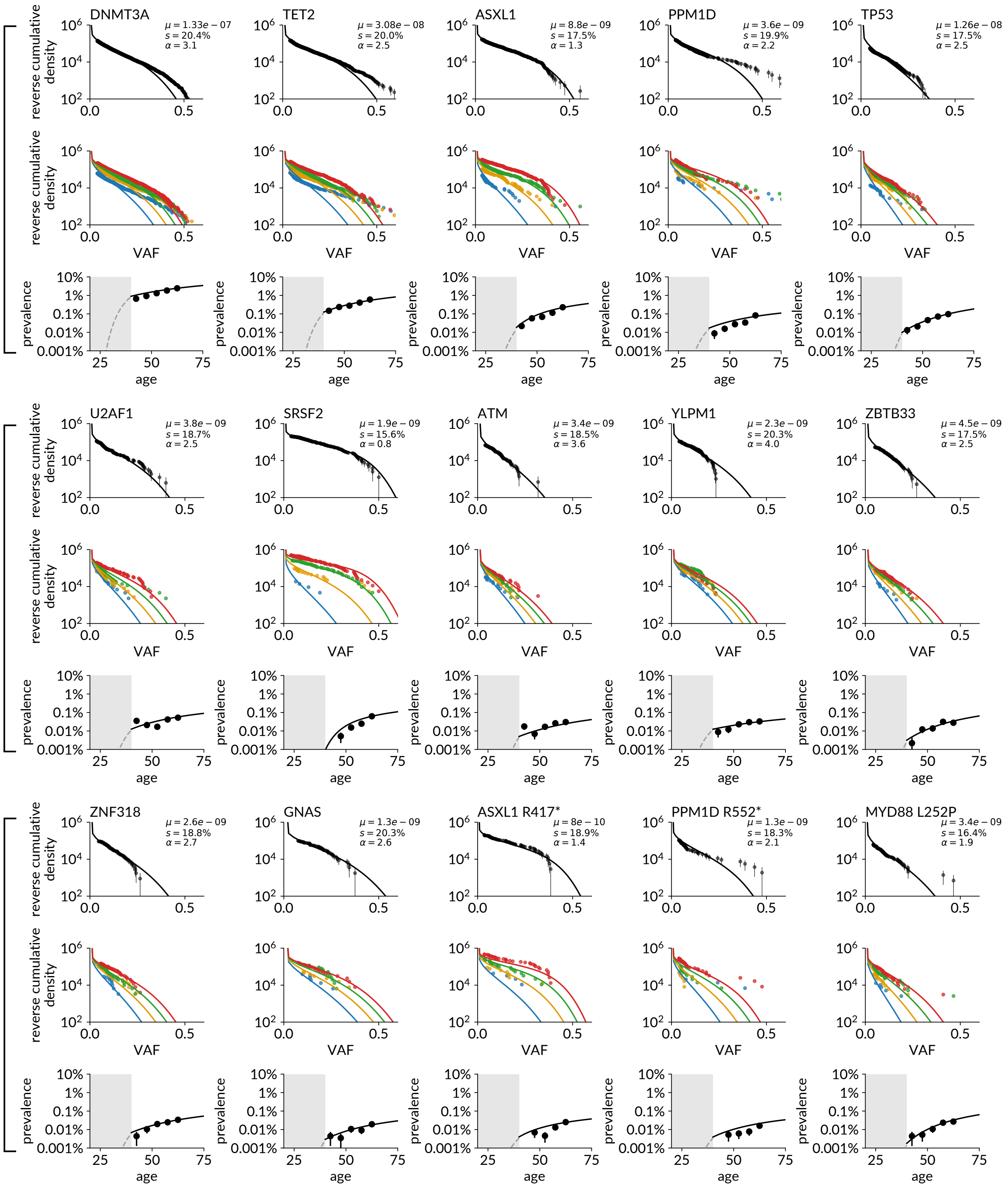


**Fig. S17. Results of two-stage model applied to 12 genes (all variants aggregated) and three additional hotspots. Top rows in each group:** Reverse cumulative VAF density for each variant, aggregated across all ages (points) compared with predictions from two-stage model. **Middle rows:** As above, stratified into four age groups. **Bottom rows:** Prevalence of each CH variant by age, compared with predicted prevalence from two-stage model. Grey shaded area shows unobserved period before age 40.

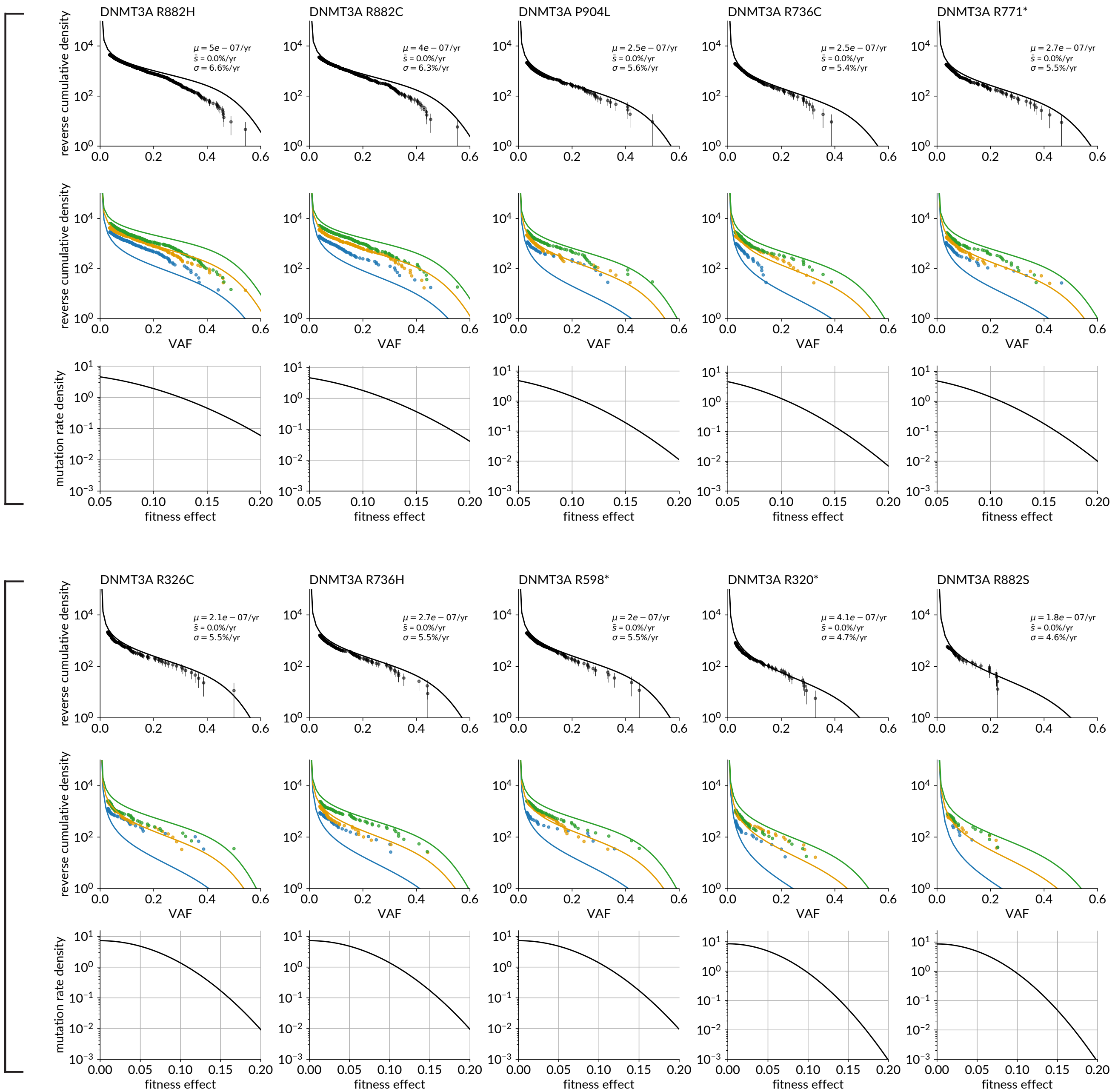


**Fig. S18. Results of distribution of fitness effects (DFE) model applied to decelerating hotspots. Top rows in each group:** Reverse cumulative VAF density for each variant, aggregated across all ages (points) compared with predictions from DFE model. **Middle rows:** As above, stratified into three age groups. **Bottom rows:** Inferred Gaussian distribution of fitness effects.

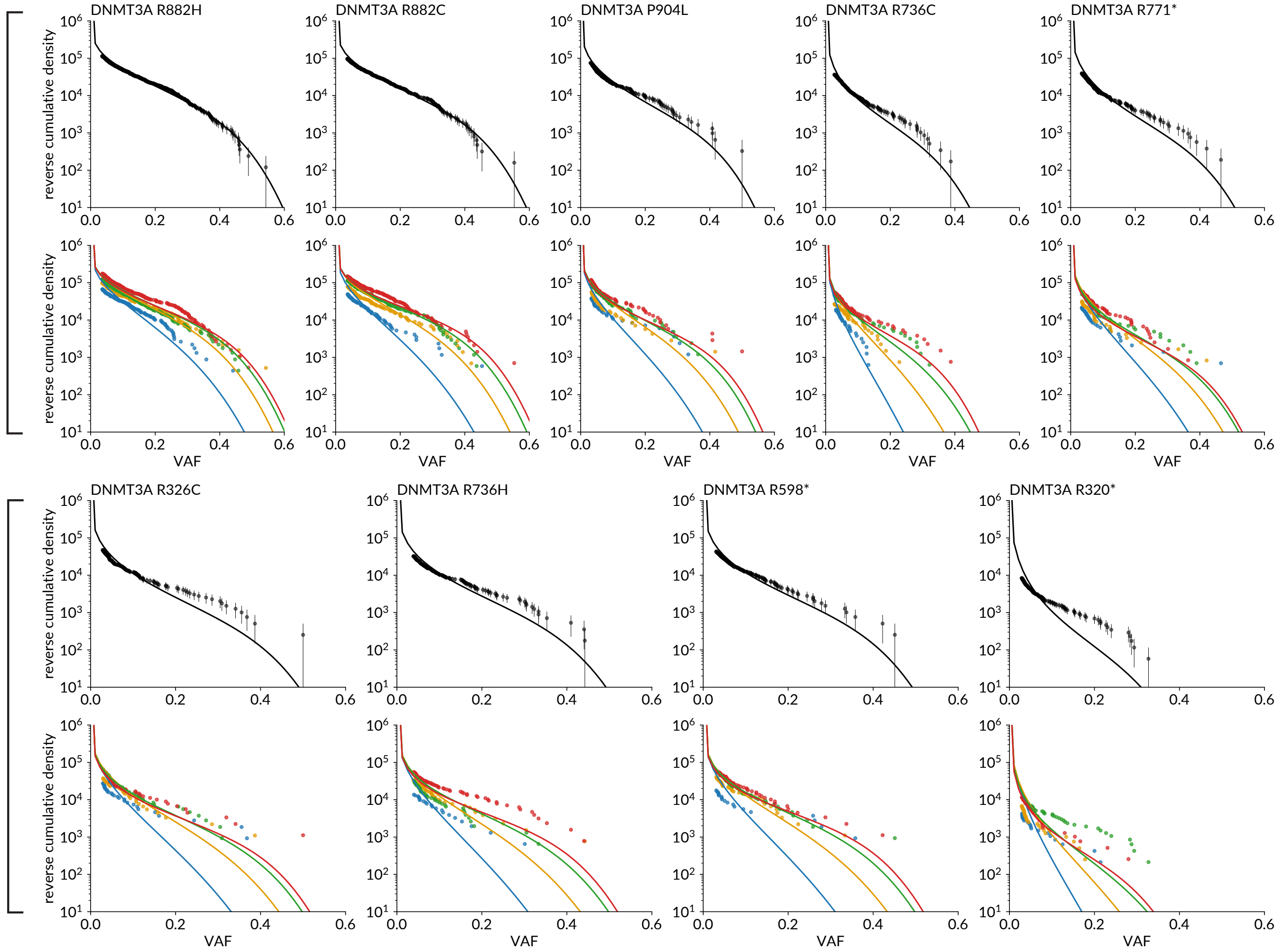


**Fig. S19. Results of clonal competition model applied to decelerating hotspots. Top rows in each group:** Reverse cumulative VAF density for each variant, aggregated across all ages (points) compared with predictions from the clonal competition model. **Bottom rows:** As above, stratified into three age groups. Inferred parameters for the model are given in Table S8.

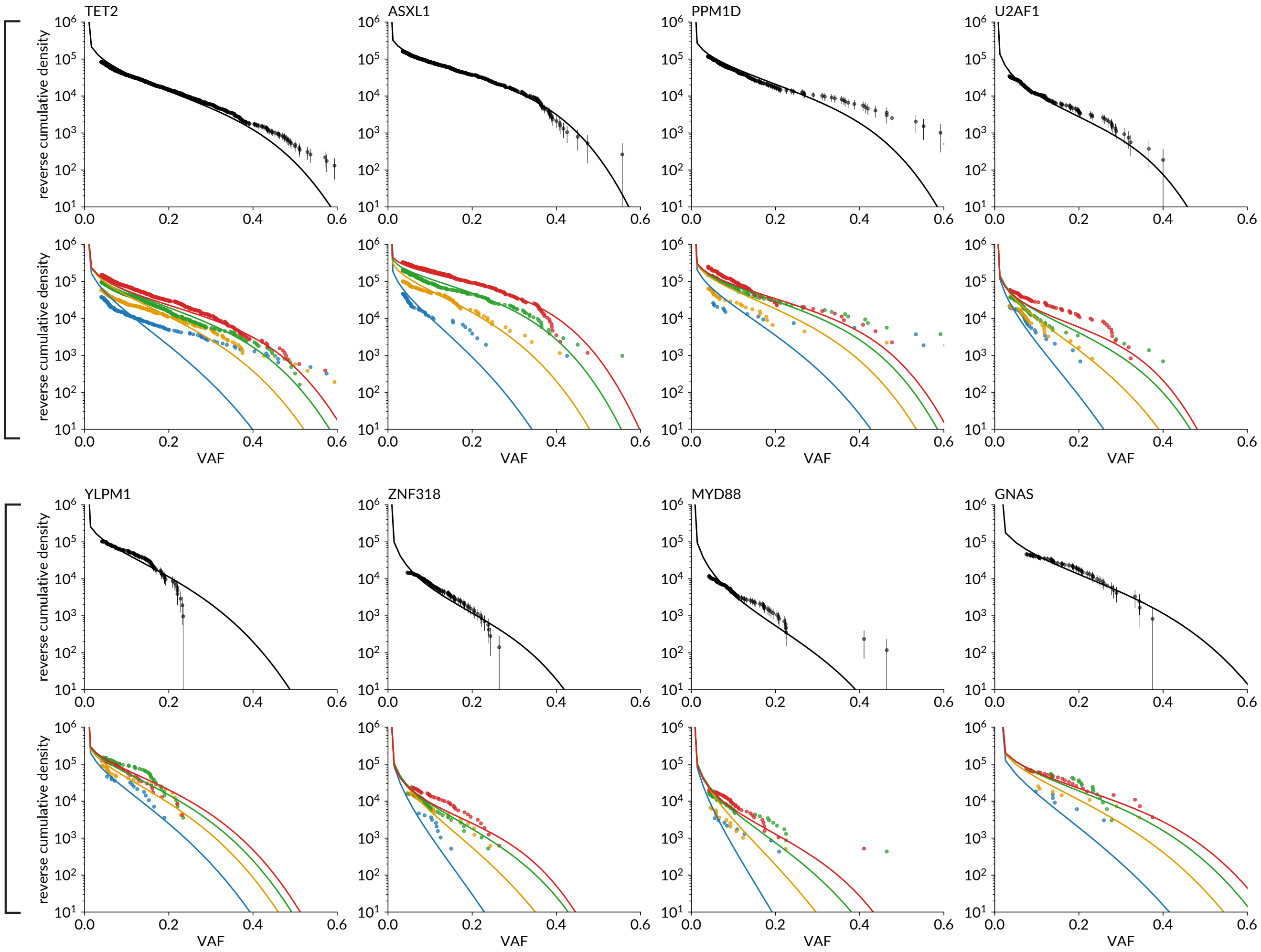


**Fig. S20. Results of clonal competition model applied to decelerating driver genes. Top rows in each group:** Reverse cumulative VAF density for each gene (variants aggregated across each gene), aggregated across all ages (points) compared with predictions from the clonal competition model. **Bottom rows:** As above, stratified into three age groups. Inferred parameters for the model are given in Table S8.

### Supplementary Note 5: Supplementary Tables


- Table S1: Genomic positions of CH hotspots
- Table S2: Constant-growth parameters for hotspots (least-squares method)
- Table S3: Constant-growth parameters for hotspots (MLE method)
- Table S4: Constant-growth parameters for exome-wide calling (MLE method)
- Table S5: Estimated parameters for two-stage model (hotspots)
- Table S6: Estimated parameters for two-stage model (exome-wide calling)
- Table S7: Estimated parameters for distribution of fitness effects model
- Table S8: Estimated parameters for clonal competition model


**Table S1. Genomic positions of CH hotspots.** Putative CH drivers targeted in the analysis, originally identified by Watson et al.[10]. Genomic positions refer to assembly build 38 (GRCh38).

| Gene | Target variant(s) | chr:position |
|---|---|---|
| *DNMT3A* | R320* | 2:25247647 |
| | R326C/G/S | 2:25247629 |
| | R598* | 2:25244214 |
| | R729G/W | 2:25240439 |
| | Y735C/F | 2:25240420 |
| | R736H/L | 2:25240417 |
| | R736C/G/S | 2:25240418 |
| | R771* | 2:25240313 |
| | R882C/S | 2:25234374 |
| | R882H/L/P | 2:25234373 |
| | W860R | 2:25235727 |
| | P904L/Q/R | 2:25234307 |
| *GNB1* | K57E | 1:1815790 |
| *IDH1* | R132H | 2:208248388 |
| *IDH2* | R140Q | 15:90088702 |
| | R172K | 15:90088606 |
| *JAK2* | V617F | 9:5073770 |
| *KIT* | D816H/Y | 4:54733154 |
| | D816V/F | 4:54733155 |
| *KRAS* | G12D/V | 12:25245350 |
| | G12C | 12:25245351 |
| *MPL* | W515L | 1:43349338 |
| *NRAS* | G12D/V | 1:114716126 |
| | G12C | 1:114716127 |
| *SF3B1* | K666N | 2:197402635 |
| | K700E | 2:197402110 |
| *SRSF2* | P95H/R/L | 17:76736877 |

**Table S2. Constant-growth model applied to CH hotspots, least-squares fitting.** Number of carriers ($n$), and inferred mutation rates ($\mu$) and fitness effects ($s$) for 18 CH hotspots using a constant-growth model, fitted using least-squares minimisation.

| Variant | $n$ | $\mu$ / per cell per year | $s$ / per year |
|---|---|---|---|
| DNMT3A R882H | 949 | $1.4 \times 10^{-8}$ | 0.155 |
| DNMT3A R882C | 606 | $8.6 \times 10^{-9}$ | 0.155 |
| DNMT3A Y735C | 373 | $7.9 \times 10^{-9}$ | 0.129 |
| DNMT3A P904L | 229 | $2.5 \times 10^{-9}$ | 0.155 |
| DNMT3A R736C | 211 | $3.5 \times 10^{-9}$ | 0.143 |
| DNMT3A R771* | 207 | $2.7 \times 10^{-9}$ | 0.151 |
| DNMT3A R326C | 187 | $1.9 \times 10^{-9}$ | 0.155 |
| DNMT3A R736H | 183 | $2.5 \times 10^{-9}$ | 0.153 |
| DNMT3A R598* | 170 | $2.2 \times 10^{-9}$ | 0.150 |
| DNMT3A R320* | 145 | $2.4 \times 10^{-9}$ | 0.140 |
| DNMT3A R729W | 127 | $1.8 \times 10^{-9}$ | 0.149 |
| SRSF2 P95H | 112 | $1.4 \times 10^{-9}$ | 0.169 |
| SF3B1 K700E | 106 | $1.1 \times 10^{-9}$ | 0.165 |
| SRSF2 P95R | 64 | $6.4 \times 10^{-10}$ | 0.171 |
| SRSF2 P95L | 51 | $4.2 \times 10^{-10}$ | 0.187 |
| DNMT3A R882S | 45 | $1.1 \times 10^{-9}$ | 0.137 |
| IDH2 R140Q | 43 | $1.6 \times 10^{-9}$ | 0.146 |
| SF3B1 K666N | 43 | $6.0 \times 10^{-10}$ | 0.171 |

**Table S3. Constant-growth model applied to CH hotspots, maximum-likelihood fitting.** Number of carriers ($n$), imposed lower VAF thresholds, and inferred mutation rates ($\mu$) and fitness effects ($s$) for 18 CH hotspots using a constant-growth model, fitted using maximum-likelihood methods.

| Variant | $n$ | VAF threshold | $\hat{\mu}$ / per cell per year | $\hat{s}$ / per year | $\log\mathcal{L}(\hat{\mu}, \hat{s})$ |
|---|---|---|---|---|---|
| DNMT3A R882H | 949 | 0.036 | $1.0 \times 10^{-8}$ | 0.175 | $-9481$ |
| DNMT3A R882C | 606 | 0.036 | $7.3 \times 10^{-9}$ | 0.169 | $-6275$ |
| DNMT3A Y735C | 373 | 0.020 | $9.0 \times 10^{-9}$ | 0.132 | $-3811$ |
| DNMT3A P904L | 229 | 0.031 | $3.0 \times 10^{-9}$ | 0.162 | $-2596$ |
| DNMT3A R736C | 211 | 0.027 | $3.5 \times 10^{-9}$ | 0.150 | $-2343$ |
| DNMT3A R771* | 207 | 0.034 | $2.7 \times 10^{-9}$ | 0.164 | $-2352$ |
| DNMT3A R326C | 187 | 0.028 | $2.4 \times 10^{-9}$ | 0.161 | $-2153$ |
| DNMT3A R736H | 183 | 0.039 | $2.8 \times 10^{-9}$ | 0.161 | $-2098$ |
| DNMT3A R598* | 170 | 0.031 | $2.4 \times 10^{-9}$ | 0.159 | $-1965$ |
| DNMT3A R320* | 145 | 0.029 | $2.5 \times 10^{-9}$ | 0.149 | $-1675$ |
| DNMT3A R729W | 127 | 0.038 | $3.1 \times 10^{-9}$ | 0.145 | $-1465$ |
| SRSF2 P95H | 112 | 0.043 | $1.8 \times 10^{-9}$ | 0.162 | $-1337$ |
| SF3B1 K700E | 106 | 0.046 | $3.5 \times 10^{-9}$ | 0.140 | $-1198$ |
| SRSF2 P95R | 64 | 0.030 | $8.4 \times 10^{-10}$ | 0.161 | $-804$ |
| SRSF2 P95L | 51 | 0.043 | $6.0 \times 10^{-10}$ | 0.174 | $-660$ |
| DNMT3A R882S | 45 | 0.037 | $9.9 \times 10^{-10}$ | 0.147 | $-560$ |
| IDH2 R140Q | 43 | 0.063 | $1.2 \times 10^{-9}$ | 0.152 | $-525$ |
| SF3B1 K666N | 43 | 0.069 | $1.2 \times 10^{-9}$ | 0.153 | $-519$ |

**Table S4. Constant-growth model applied to exome-wide CH calls, maximum-likelihood fitting.** Number of carriers ($n$), imposed lower VAF thresholds, and inferred mutation rates ($\mu$) and fitness effects ($s$) inferred for 13 CH driver genes (aggregated) and 3 additional hotspots from the exome-wide CH calls, using a constant-growth model, fitted using maximum-likelihood methods.

| Gene | n | VAF threshold | $\hat{\mu}$ / per cell per year | $\hat{s}$ / per year |
|---|---|---|---|---|
| DNMT3A | 7799 | 0.040 | $1.1 \times 10^{-7}$ | 0.165 |
| TET2 | 1895 | 0.040 | $2.6 \times 10^{-8}$ | 0.165 |
| ASXL1 | 624 | 0.034 | $8.7 \times 10^{-9}$ | 0.162 |
| TP53 | 294 | 0.048 | $8.2 \times 10^{-9}$ | 0.145 |
| PPM1D | 234 | 0.039 | $3.2 \times 10^{-9}$ | 0.165 |
| U2AF1 | 183 | 0.034 | $3.2 \times 10^{-9}$ | 0.153 |
| SF3B1 | 172 | 0.059 | $5.1 \times 10^{-9}$ | 0.148 |
| SRSF2 | 166 | 0.037 | $1.9 \times 10^{-9}$ | 0.173 |
| CBL | 128 | 0.060 | $3.2 \times 10^{-9}$ | 0.153 |
| YLPM1 | 107 | 0.041 | $1.7 \times 10^{-9}$ | 0.161 |
| ZNF318 | 104 | 0.042 | $2.0 \times 10^{-9}$ | 0.154 |
| ZBTB33 | 101 | 0.050 | $2.9 \times 10^{-9}$ | 0.145 |
| MYD88 | 101 | 0.042 | $2.6 \times 10^{-9}$ | 0.145 |
| ATM | 99 | 0.040 | $2.1 \times 10^{-9}$ | 0.150 |
| GNAS | 56 | 0.071 | $9.8 \times 10^{-10}$ | 0.169 |

**Table S5. Two-stage model applied to CH hotspots.** Mutation rates ($\mu$), initial fitness effects ($s_1$) and deceleration factors ($\alpha$) inferred for 18 CH hotspots using a two-stage evolutionary model, fitted using maximum-likelihood methods. Test statistic and p-values derive from likelihood ratio test with a constant-growth model ($s = s_1$, $\alpha = 1$), using a $\chi^2$ distribution with one degree of freedom.

| Variant | $n$ | VAF threshold | $\hat{\mu}$ / per cell per year | $\hat{s_1}$ / per year | $\hat{\alpha}$ | 95% CI | $\chi^2(1)$ | $p$ |
|---|---|---|---|---|---|---|---|---|
| DNMT3A R882H | 949 | 0.036 | $1.3 \times 10^{-8}$ | 0.219 | 4.1 | 3.3-5.4 | 359.5 | $< 1 \times 10^{-15}$ |
| DNMT3A R882C | 606 | 0.036 | $8.8 \times 10^{-9}$ | 0.209 | 3.1 | 2.5-4.1 | 178.7 | $< 1 \times 10^{-15}$ |
| DNMT3A Y735C | 373 | 0.020 | $1.0 \times 10^{-8}$ | 0.157 | 2.0 | 1.6-2.6 | 46.6 | $8.5 \times 10^{-12}$ |
| DNMT3A P904L | 229 | 0.031 | $3.7 \times 10^{-9}$ | 0.202 | 3.4 | 2.4-5.6 | 80.7 | $< 1 \times 10^{-15}$ |
| DNMT3A R736C | 211 | 0.027 | $4.0 \times 10^{-9}$ | 0.182 | 2.4 | 1.8-3.5 | 40.8 | $1.7 \times 10^{-10}$ |
| DNMT3A R771* | 207 | 0.034 | $3.9 \times 10^{-9}$ | 0.208 | 5.2 | 3.3-11.5 | 112.6 | $< 1 \times 10^{-15}$ |
| DNMT3A R326C | 187 | 0.028 | $3.2 \times 10^{-9}$ | 0.203 | 4.4 | 2.8-9.7 | 74.1 | $< 1 \times 10^{-15}$ |
| DNMT3A R736H | 183 | 0.039 | $3.7 \times 10^{-9}$ | 0.197 | 3.1 | 2.2-4.8 | 63.3 | $1.8 \times 10^{-15}$ |
| DNMT3A R598* | 170 | 0.031 | $3.1 \times 10^{-9}$ | 0.200 | 3.7 | 2.4-7.7 | 56.7 | $5.1 \times 10^{-14}$ |
| DNMT3A R320* | 145 | 0.029 | $3.6 \times 10^{-9}$ | 0.188 | 3.9 | 2.5-7.9 | 63.2 | $1.9 \times 10^{-15}$ |
| DNMT3A R729W | 127 | 0.038 | $4.6 \times 10^{-9}$ | 0.177 | 3.0 | 2.0-4.9 | 39.2 | $3.9 \times 10^{-10}$ |
| SRSF2 P95H | 112 | 0.043 | $1.8 \times 10^{-9}$ | 0.148 | 0.8 | 0.6-1.1 | 1.7 | 0.19 |
| SF3B1 K700E | 106 | 0.046 | $3.7 \times 10^{-9}$ | 0.153 | 1.3 | 0.9-1.8 | 2.5 | 0.11 |
| SRSF2 P95R | 64 | 0.030 | $8.5 \times 10^{-10}$ | 0.150 | 0.8 | 0.5-1.4 | 0.6 | 0.45 |
| SRSF2 P95L | 51 | 0.043 | $6.0 \times 10^{-10}$ | 0.159 | 0.8 | 0.5-1.4 | 0.8 | 0.38 |
| DNMT3A R882S | 45 | 0.037 | $1.4 \times 10^{-9}$ | 0.180 | 2.9 | 1.5-9.0 | 12.1 | 0.00051 |
| SF3B1 K666N | 43 | 0.069 | $1.1 \times 10^{-9}$ | 0.107 | 0.5 | 0.1-1.0 | 4.3 | 0.038 |
| IDH2 R140Q | 43 | 0.063 | $1.2 \times 10^{-9}$ | 0.157 | 1.1 | 0.6-2.0 | 0.2 | 0.7 |

**Table S6. Two-stage model applied to exome-wide CH calls.** Mutation rates ($\mu$), initial fitness effects ($s_1$) and deceleration factors ($\alpha$) inferred for 13 CH genes (aggregated) and three additional hotspots using a two-stage evolutionary model, fitted using maximum-likelihood methods. Test statistic and p-values derive from likelihood ratio test with a constant-growth model ($s = s_1$, $\alpha = 1$), using a $\chi^2$ distribution with one degree of freedom.

| Gene/Variant | $n$ | VAF threshold | $\hat{\mu}$ / per cell per year | $\hat{s_1}$ / per year | $\hat{\alpha}$ | 95% CI | $\chi^2(1)$ | $p$ |
|---|---|---|---|---|---|---|---|---|
| DNMT3A | 7799 | 0.040 | $1.3 \times 10^{-7}$ | 0.204 | 3.1 | $3.0 - 3.4$ | 2653.0 | $< 1 \times 10^{-15}$ |
| TET2 | 1895 | 0.040 | $3.1 \times 10^{-8}$ | 0.200 | 2.5 | 2.2-2.8 | 427.6 | $< 1 \times 10^{-15}$ |
| ASXL1 | 624 | 0.034 | $8.8 \times 10^{-9}$ | 0.175 | 1.3 | 1.1-1.5 | 13.7 | 0.00021 |
| TP53 | 294 | 0.048 | $1.3 \times 10^{-8}$ | 0.175 | 2.5 | 2.0-3.3 | 75.4 | $< 1 \times 10^{-15}$ |
| PPM1D | 234 | 0.039 | $3.6 \times 10^{-9}$ | 0.199 | 2.2 | 1.7-3.2 | 41.9 | $9.8 \times 10^{-11}$ |
| U2AF1 | 183 | 0.034 | $3.8 \times 10^{-9}$ | 0.187 | 2.5 | 1.9-3.6 | 52.0 | $5.5 \times 10^{-13}$ |
| SF3B1 | 172 | 0.059 | $5.5 \times 10^{-9}$ | 0.164 | 1.4 | 1.1-1.8 | 6.8 | 0.0093 |
| SRSF2 | 166 | 0.037 | $1.9 \times 10^{-9}$ | 0.156 | 0.8 | 0.6-1.0 | 2.9 | 0.09 |
| YLPM1 | 107 | 0.041 | $2.3 \times 10^{-9}$ | 0.203 | 4.0 | 2.4-10.1 | 44.7 | $2.3 \times 10^{-11}$ |
| ZNF318 | 104 | 0.042 | $2.7 \times 10^{-9}$ | 0.188 | 2.7 | 1.8-4.8 | 29.5 | $5.7 \times 10^{-8}$ |
| MYD88 | 101 | 0.042 | $3.3 \times 10^{-9}$ | 0.174 | 2.2 | 1.5-3.5 | 20.9 | $5 \times 10^{-6}$ |
| ZBTB33 | 101 | 0.050 | $4.5 \times 10^{-9}$ | 0.175 | 2.5 | 1.7-4.1 | 24.5 | $7.4 \times 10^{-7}$ |
| ATM | 99 | 0.040 | $3.4 \times 10^{-9}$ | 0.185 | 3.6 | 2.2-7.1 | 42.1 | $8.6 \times 10^{-11}$ |
| MYD88 L252P | 89 | 0.042 | $3.4 \times 10^{-9}$ | 0.164 | 1.9 | 1.3-2.9 | 12.2 | 0.00048 |
| ASXL1 R417* | 70 | 0.034 | $8.1 \times 10^{-10}$ | 0.189 | 1.4 | 1.0-2.4 | 2.9 | 0.09 |
| PPM1D R552* | 58 | 0.039 | $1.3 \times 10^{-9}$ | 0.183 | 2.1 | 1.4-3.9 | 13.0 | 0.00031 |
| GNAS | 56 | 0.071 | $1.3 \times 10^{-9}$ | 0.203 | 2.6 | 1.5-5.8 | 15.0 | 0.00011 |

**Table S7. Distribution of fitness effects model.** Mutation rates ($\mu$) and distribution of fitness effects inferred for 12 decelerating CH driver variants using a Gaussian distribution of fitness effects with mean fitness $\bar{s}$ and standard deviation $\sigma$, fitted using maximum-likelihood methods.

| Variant | $n$ | $\hat{\mu}$ / per cell per year | $\hat{\bar{s}}$ / per year | $\hat{\sigma}$ / per year | $\chi^2(1)$ | $p$ |
|---|---|---|---|---|---|---|
| DNMT3A R882H | 949 | $5.0 \times 10^{-7}$ | 0.000 | 0.066 | 354.9 | $< 1 \times 10^{-15}$ |
| DNMT3A R882C | 606 | $4.0 \times 10^{-7}$ | 0.000 | 0.063 | 227.7 | $< 1 \times 10^{-15}$ |
| DNMT3A P904L | 229 | $2.5 \times 10^{-7}$ | 0.000 | 0.056 | 147.3 | $< 1 \times 10^{-15}$ |
| DNMT3A R736C | 211 | $2.5 \times 10^{-7}$ | 0.000 | 0.054 | 48.9 | $2.6 \times 10^{-12}$ |
| DNMT3A R771* | 207 | $2.7 \times 10^{-7}$ | 0.000 | 0.055 | 141.9 | $< 1 \times 10^{-15}$ |
| DNMT3A R326C | 187 | $2.1 \times 10^{-7}$ | 0.000 | 0.055 | 116.3 | $< 1 \times 10^{-15}$ |
| DNMT3A R736H | 183 | $2.7 \times 10^{-7}$ | 0.000 | 0.055 | 87.8 | $< 1 \times 10^{-15}$ |
| DNMT3A R598* | 170 | $2.0 \times 10^{-7}$ | 0.000 | 0.055 | 87.2 | $< 1 \times 10^{-15}$ |
| DNMT3A R320* | 145 | $4.1 \times 10^{-7}$ | 0.000 | 0.047 | 81.2 | $< 1 \times 10^{-15}$ |
| DNMT3A R882S | 45 | $1.8 \times 10^{-7}$ | 0.000 | 0.046 | 14.3 | 0.00016 |

Model did not converge for DNMT3A Y735C and DNMT3A R729W

**Table S8. Clonal competition model.** Mutation rates ($\mu$) and distribution of fitness effects ($s$) for decelerating genes and DNMT3A hotspots. Parameters $\mu_1, s_1$ are for the observed driver variant, $\mu_2, s_2$ for the hidden background clones.

| Gene/Variant | $n$ | $\hat{\mu}_1$ / per cell per year (variant) | $\hat{s}_1$ / per year (variant) | $\hat{\mu}_2$ / per cell per year (background) | $\hat{s}_2$ / per year (background) |
|---|---|---|---|---|---|
| DNMT3A R882H | 949 | $2.0 \times 10^{-8}$ | 0.204 | $1.1 \times 10^{-5}$ | 0.228 |
| DNMT3A R882C | 606 | $1.5 \times 10^{-8}$ | 0.190 | $8.4 \times 10^{-6}$ | 0.224 |
| DNMT3A P904L | 229 | $7.2 \times 10^{-9}$ | 0.185 | $1.0 \times 10^{-5}$ | 0.226 |
| DNMT3A R736C | 211 | $1.4 \times 10^{-8}$ | 0.157 | $7.3 \times 10^{-6}$ | 0.225 |
| DNMT3A R771* | 207 | $1.2 \times 10^{-8}$ | 0.186 | $1.1 \times 10^{-5}$ | 0.238 |
| DNMT3A R326C | 187 | $9.3 \times 10^{-9}$ | 0.180 | $1.1 \times 10^{-5}$ | 0.232 |
| DNMT3A R736H | 183 | $1.3 \times 10^{-8}$ | 0.172 | $9.0 \times 10^{-6}$ | 0.230 |
| DNMT3A R598* | 170 | $9.3 \times 10^{-9}$ | 0.173 | $9.0 \times 10^{-6}$ | 0.230 |
| DNMT3A R320* | 145 | $4.1 \times 10^{-8}$ | 0.150 | $1.1 \times 10^{-5}$ | 0.229 |
| TET2 | 1895 | $5.4 \times 10^{-8}$ | 0.182 | $9.1 \times 10^{-6}$ | 0.213 |
| ASXL1 | 624 | $9.0 \times 10^{-9}$ | 0.170 | $3.7 \times 10^{-5}$ | 0.128 |
| PPM1D | 234 | $4.7 \times 10^{-9}$ | 0.190 | $1.2 \times 10^{-5}$ | 0.203 |
| U2AF1 | 183 | $1.3 \times 10^{-8}$ | 0.160 | $6.3 \times 10^{-6}$ | 0.230 |
| YLPM1 | 107 | $2.4 \times 10^{-9}$ | 0.206 | $1.2 \times 10^{-4}$ | 0.177 |
| ZNF318 | 104 | $1.7 \times 10^{-8}$ | 0.155 | $7.3 \times 10^{-6}$ | 0.229 |
| MYD88 | 101 | $2.0 \times 10^{-8}$ | 0.147 | $9.4 \times 10^{-6}$ | 0.213 |
| GNAS | 56 | $2.9 \times 10^{-9}$ | 0.179 | $1.1 \times 10^{-5}$ | 0.204 |

Model did not converge for DNMT3A Y735C, DNMT3A R729W, DNMT3A R882S, TP53, ZBTB33, ATM
